# Stretch-induced endogenous electric fields drive neural crest directed collective cell migration *in vivo*

**DOI:** 10.1101/2021.10.11.463916

**Authors:** Fernando Ferreira, Sofia Moreira, Elias H. Barriga

## Abstract

Directed collective cell migration (dCCM) is essential for morphogenesis^1, 2^. Cell clusters migrate in inherently complex *in vivo* environments composed of chemical, electrical, mechanical as well as topological features. While these environmental factors have been shown to allow dCCM *in vitro*, our understanding of dCCM *in vivo* is mostly limited to chemical guidance^3^. Thus, despite its wide biological relevance, the mechanisms that guide dCCM *in vivo* remain unclear. To address this, we study endogenous electric fields in relation to the migratory environment of the *Xenopus laevis* cephalic neural crest, an embryonic cell population that collectively and directionally migrates *in vivo*^4^, and whose migratory mode has been linked to cancer invasion and metastasis^5^. Combining bioelectrical, biomechanical and molecular tools, we show that endogenous electric fields drive neural crest dCCM via electrotaxis *in vivo*. Moreover, we identify the voltage-sensitive phosphatase 1 (Vsp1) as a key component of the molecular mechanism used by neural crest cells to transduce electric fields into a directional cue *in vivo*. Furthermore, Vsp1 function is specifically required for electrotaxis, being dispensable for cell motility and chemotaxis. Finally, we reveal that endogenous electric fields are mechanoelectrically established. Mechanistically, convergent extension movements of the neural fold generate membrane tension, which in turn opens stretch-activated channels to mobilise the ions required to fuel electric fields. Overall, our results reveal a mechanism of cell guidance, where electrotaxis emerges from the mechanoelectrical and molecular interplay between neighbouring tissues. More broadly, our data contribute to validate the, otherwise understudied, functions of endogenous bioelectrical stimuli in morphogenetic processes^6^.

## Main

Cephalic neural crest cells are induced at the interface between the neural plate and the non-neural ectoderm, from where they migrate following stereotypical mediolateral paths known as streams^7^ (**Fig. 1a**). The migration of neural crest cells has been widely studied; yet, how do they attain directionality in convoluted native contexts remains unclear. Chemical cues have been proposed to guide the directed collective cell migration (dCCM) of neural crest cells via chemotaxis^8^. However, recent evidence suggested that chemotaxis may not be sufficient to explain their directed motion *in vivo*^9^. Intriguingly, *in vitro* experiments have demonstrated the ability of neural crest cells to directionally migrate by following extrinsically imposed electric fields, in a process termed as electrotaxis (or galvanotaxis)^10–12^. In addition, seminal work detected the presence of electric fields in amphibian embryos during neurulation^13, 14^ and the application of external electric fields led to developmental defects^15^. In this scenario, electrotaxis emerges as an alternative or complementary mechanism to explain neural crest dCCM *in vivo*. Nonetheless, whether and how endogenous electric fields drive directed cell migration via electrotaxis in native contexts remains unclear.

**Fig. 1.**
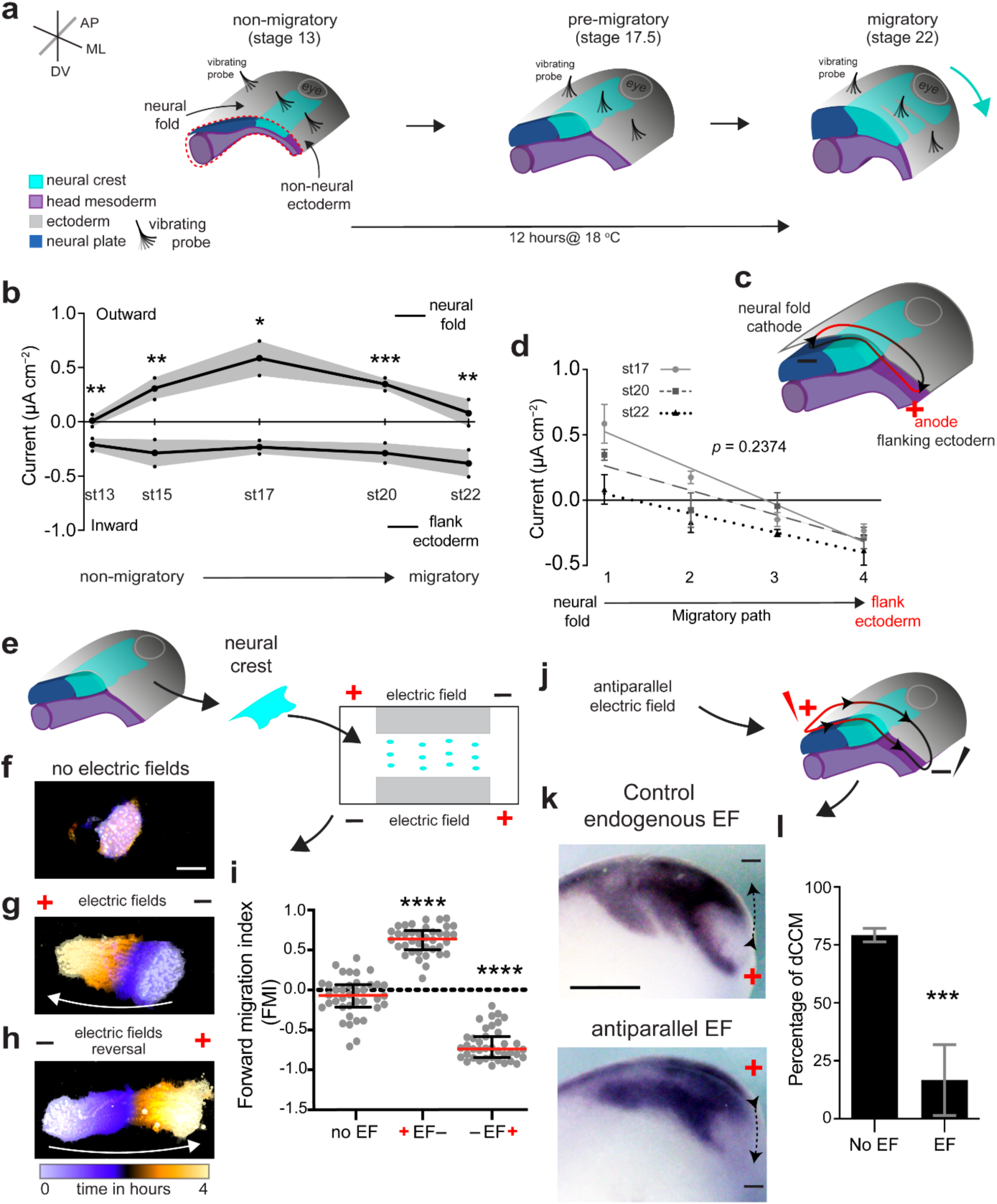
**Endogenous electric fields modulate neural crest directed collective cell migration. a**, Scheme shows neural crest development and vibrating probe measurements along its migratory path (AP, anteroposterior; ML, mediolateral; DV, dorsoventral). **b–d**, Mediolateral endogenous electric fields (EFs) emerge in the migratory path of neural crest cells. **b**, Vibrating probe measurements results; dots represent mean and shade the standard error, paired *t*-test (st13–20) or Wilcoxon test (st22) (both two-tailed), \*\**p*_st13_ = 0.0017, \*\**p*_st15_ = 0.0042, \**p*_st17_ = 0.0339, \*\*\**p*_st20_ = 0.0008, \*\**p*_st22_ = 0.0078, *n* = 35 embryos. **c**, Sustained EF dipole along the migratory path of neural crest cells, the cathode (−) is in the neural folds and the anode (+) in the flanking non-neural ectoderm. **d**, Linear regressions showing similar slopes of current along the neural crest migratory path at different stages, *p* = 0.2374 (two-tailed), *n* = 23 embryos. **e– i**, Neural crest explants exposed to EFs *ex vivo*. **e**, *Ex vivo* electrotaxis assay diagram. **f–h**, Time colour-coded trajectories of neural crest cells migrating in the absence of EFs for 1 h (**f**), under EFs (100 mV mm^−1^) for 4 h (**g**) or after EFs reversal for 4 h (**h**). White arrows indicate direction of neural crest migration (see **Supplementary Video 1**). Scale bar, 100 μm. **i**, Forward migration index (FMI) quantifications, conditions as indicated. Red lines represent median and error bars the interquartile ranges, paired *t-*test, \*\*\*\**p* < 0.0001 (two-tailed), *n* = 39 clusters. **j– l**, Antiparallel EFs *in vivo* alter the pattern of neural crest dCCM. **j**, Antiparallel EFs application (100 mV mm^−1^). **k**, *In situ* hybridisations showing lateral views of embryos hybridised with *c*3 (a marker of neural crest migration). Scale bar, 200 μm. **l**, Percentage of embryos displaying streams, column bars represent mean and error bars the standard deviation, Fisher’s exact test, \*\*\**p* = 0.0002 (two-tailed), *n*_No EFs_ = 30 embryos, *n*_Antiparallel EFs_ = 18 embryos. **f–h**,**k**, Representative examples from at least three independent experiments; CI = 95%.

To address this, we first used ultrasensitive vibrating probes to map extracellular electric currents^16^ along the migratory path of neural crest cells. We performed our measurements from the neural fold towards the flanking non-neural ectoderm and from non-migratory (stage 13) to migratory (stage 22) stages of neural crest cells (**Fig. 1a; Supplementary Fig.. 1**). Our spatiotemporal measurements detected a long-lasting and sustained dipole loop, with outward currents registered in the neural fold and inward currents in the flanking non-neural ectoderm (**Fig. 1b**). Thus, based in Ohm’s law^17^ (**Methods**), this pattern of electric currents indicates that a mediolateral endogenous electric field emerges in the neural crest migratory path from early stages of migration, with the cathode (− pole) in the neural fold and the anode (+ pole) in the flanking non-neural ectoderm (**Fig. 1c**). Furthermore, we observed that outward currents reached a maximum at stages in which neural crest cells are preparing to migrate (pre-migratory stages 17 and 20) and that these values decreased once migration started (**Fig. 1b**). Despite these differences in the absolute magnitude of current, the slope of the electric field established in the migratory path of neural crest cells remains unaltered from pre-migratory stages (**Fig. 1d**). These results indicate that neural crest cells may experience electric fields towards the onset of their dCCM.

To dissect the role of these endogenous electric fields in neural crest directed cell migration, we generated an *ex vivo* electrotaxis assay capable of reproducing the recorded endogenous electric fields in a controlled microenvironment (**Fig. 1e; Methods**). In the absence of an electric field, cultured neural crests clusters tend to radially disperse and to perform random migration (**Supplementary Video 1**), but when an electric field was applied, the collectiveness of these clusters is retained and the groups of neural crest cells persistently and collectively migrated towards the anode (**Fig. 1f,g**; **Supplementary Video 1**). Interestingly, switching the polarity of the electric field led to a switch in the direction of migration towards the new anodal position (**Fig. 1h**; **Supplementary Video 1**). Note that this direction of migration agrees with the orientation of the endogenous electric field recorded *in vivo* (**Fig. 1d**). These observations were quantitatively confirmed by computing the forward migration index (FMI) as a readout of the directionality displayed by clusters in each condition (**Fig. 1i**); while FMIs closer to 0 represent random migration, FMIs near 1 or −1 account for a directional response (**Methods**). Since the recorded endogenous electric fields ranged from ∼10–100 mV mm^−1^, we tested the response of neural crest cells to electric fields of 10, 25 and 100 mV mm^−1^ (**Methods**). Our assays confirmed that the directionality of neural crest cells was consistent in the range of electric fields analysed and that, until some extent, cluster directionality depends on the level of the applied electric fields (**Supplementary Fig.. 2; Supplementary Video 2**). Next, we addressed whether the directional response of neural crest cells to electric fields is an emergent property of the collective or also appears in individual cells. By comparing the response of neural crest clusters with the behaviour of single cells exposed to the same electric field, we found that isolated cells displayed FMIs closer to 0, whereas clusters displayed effective anodal response (**Supplementary Fig.. 3; Supplementary Video 3**). Together, these results indicate that physiological levels of electric fields influence the directionality of neural crest cells and consequently their collective migration *ex vivo*. Thus, to confirm that these electric fields are relevant for neural crest migration *in vivo*, we perturbed the endogenous electric fields by extrinsically applying antiparallel electric fields to the neural crest migratory path (**Fig. 1j; Methods**). While control neural crests migrated by following stereotypical streams (a hallmark of dCCM^18, 19^), the antiparallel electric field consistently impaired neural crest streams formation, as shown by *in situ* hybridisation of a migratory neural crest marker (**Fig. 1k,l**). Altogether, our data indicate that endogenous electric fields are required for the dCCM of neural crest cells *in vivo*, introducing electrotaxis as a complementary or alternative mechanism by which the neural crest attain their characteristic directionality.

To explore this possibility and to gain insights into the mechanism of electrotaxis *in vivo*, we investigated how neural crest cells sense and transduce endogenous electric fields into a directional cue. Most of the proteins reported to mediate electrotaxis are part of the cell self-polarity^20, 21^ or cell-substrate adhesion^22, 23^ machineries and few molecules have been shown to specifically mediate electrotaxis *in vitro*^24–27^. Hence, the precise mechanism used by cells to transduce electric fields into a directional response *in vivo* remains unclear. To address this point, we searched for candidate molecules for the mediation of electrotaxis (electrosensors) in the neural crest by performing RNA-seq experiments from isolated neural crest clusters (**Supplementary Fig.. 4**). Among other candidates, we found that the voltage-sensitive phosphatase 1 (originally described as *tpte*2*.L* and referred here as *vsp*1)^28, 29^ was enriched in our neural crest-specific library, whereas the paralogous gene *vsp*2 (*tpte*2*.S*) was not expressed (**Supplementary Table. 1; Supplementary Fig.. 4**). Vsp1 contains a channel-like transmembrane domain that senses voltage fluctuations and a cytosolic catalytic domain with phosphatase activity^28, 30^. This peculiar protein structure positioned Vsp1 as a good candidate to sense and transduce electrical stimuli into a cascade that could eventually lead to electrotaxis. To test this, we first validated the expression of Vsp1 in neural crest cells and confirmed its localization to the cell membrane, where it is expected to operate upon membrane depolarization (**Supplementary Fig.. 4**). Then, we functionally tested the requirement of Vsp1 activity for neural crest electrotaxis *ex vivo*, by expressing a catalytically inactive form of Vsp1 (Vsp1-C301S) previously shown to reduce the phosphatase activity in *Xenopus* oocytes to applied voltage^29, 31^. In the absence of electric fields, Vsp1-C301S injected neural crest cells did not display any significant effects in motility, as observed in our *ex vivo* assays (**Supplementary Fig.. 5**; **Supplementary Video 4**). However, when an electric field was applied, Vsp1-C301S injected clusters failed to directionally respond, unlike control clusters that effectively performed electrotaxis (**Fig. 2a–c**; **Supplementary Video 5**), strongly suggesting a role for Vsp1 in collective electrotaxis. To further test the involvement of Vsp1 in electrotaxis, we investigated whether Vsp1 activity is sufficient to allow directed migration at suboptimal levels of electric fields. First, we determined that electrotaxis of wild-type neural crest clusters was minimal when exposed to electric fields of 5 mV mm^−1^ *ex vivo* (**Fig. 2d**). Then, we injected a mutant form of Vsp1 which is more sensitive to voltage (VSP1-R152Q-GFP)^29, 31^ and analysed its impact in neural crest electrotaxis. VSP1-R152Q-GFP expression was sufficient to significantly increase the frequency of neural crest clusters displaying a persistent and directional response under these suboptimal electric fields (**Fig. 2d–f**; **Supplementary Video 6**), confirming that Vsp1 activity modulates the sensitivity of cells to electric fields.

**Fig. 2.**
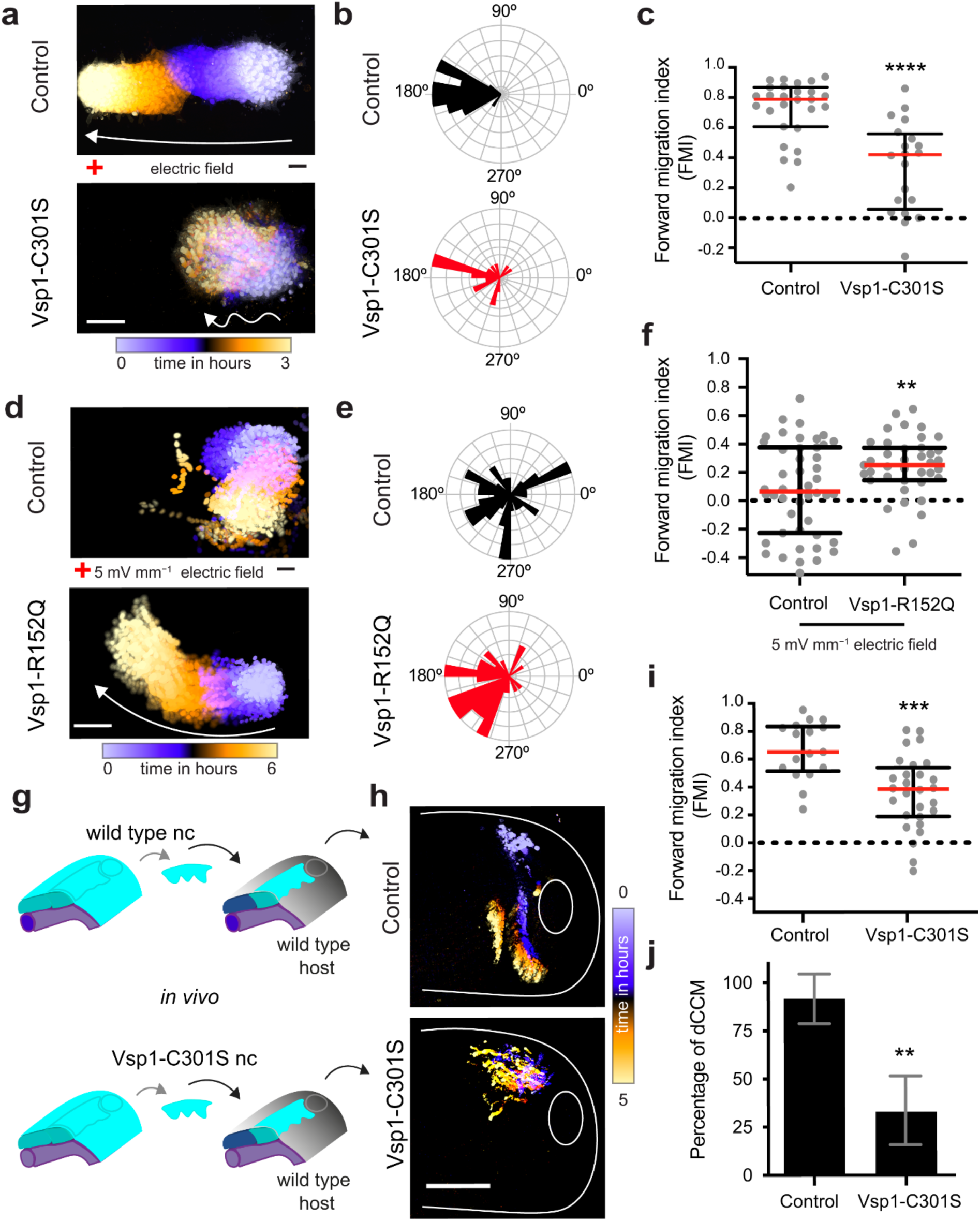
**Voltage-sensitive phosphatase 1 (Vsp1) controls neural crest directionality. a–c**, Effect of Vsp1 (Vsp1-C301S) in neural crest dCCM *ex vivo*. **a**, Time colour-coded trajectories of neural crest cells *ex vivo*, white arrows indicate direction of neural crest migration (see **Supplementary Video 5**). Scale bar, 100 μm. **b**, Rose plots showing the angle frequencies of neural crest migration in relation to the electric field (EF) vector (anode at 180°). **c**, Forward migration index (FMI) quantifications, red lines represent median and error bars the interquartile ranges, Mann Whitney *U*-test, \*\*\*\**p* < 0.0001 (two-tailed), *n*_Control_ = 27 clusters, *n*_Vsp1-C301S_ = 20 clusters. **d–f**, Effect of Vsp1 (Vsp1-R152Q) in neural crest directionality exposed to suboptimal EFs *ex vivo*. **d**, Time colour-coded trajectories of neural crest cells (see **Supplementary Video 6**). Scale bar, 100 μm. **e**, Rose plots showing the angle frequencies of neural crest migration in relation to the vector of suboptimal EFs (anode at 180°). **f**, FMI quantifications, red lines represent median and error bars the interquartile ranges, *t*-test with Welch’s correction, \*\**p* = 0.0069 (two-tailed), *n*_Control_ = 47 clusters, *n*_Vsp1-R152Q_ = 39 clusters. **g–j**, Effect of Vsp1-C301S in neural crest dCCM *in vivo*. **g**, Scheme of the transplantation assays (nc, neural crest). **h**, Time colour-coded trajectories of neural crest cells *in vivo* (lateral view of embryo) (see **Supplementary Video 7**). Scale bar, 200 μm. **i**, FMI quantifications, red lines represent median and error bars the interquartile ranges, *t*-test with Welch’s correction, \*\*\**p* = 0.0002 (two-tailed), *n*_Control_ = 17 embryos, *n*_Vsp1-C301S_ = 28 embryos. **j**, Percentage of embryos displaying streams, column bars represent mean and error bars the standard deviation, Fisher’s exact test, \*\**p* = 0.0016 (two-tailed), *n*_Control_ = 17 embryos, *n*_Vsp1-C301S_ = 29 embryos. **a**,**d**,**h**, Representative examples from at least three independent experiments; CI = 95%.

Next, we tested the requirement of Vsp1 for electrotaxis *in vivo* by analysing neural crest migration in embryos injected with Vsp1-C301S. Vsp1 loss-of-function showed a consistent decrease in the formation of stereotypical streams, as shown by *in situ* hybridisation analyses (**Supplementary Fig.. 5),** suggesting an effect in directionality. Hence, we next performed graft experiments in which control or Vsp1-C301S nuclei-tagged neural crest cells were transplanted into wild-type untagged embryos. Our analyses of cell trajectories revealed that while control cells effectively migrated by forming stereotypical streams, the directionality of Vsp1-C301S injected cells was significantly reduced *in vivo* (**Fig. 2g–j**; **Supplementary Video 7**). Furthermore, since chemotaxis has been also proposed to guide neural crest migration^32^, we exposed control and Vsp1-C301S injected clusters to a previously described chemotaxis assay relying in the response of neural crest clusters to the chemokine stromal cell-derived factor 1 (SDF-1, also known as CXCL12)^32^. Our results showed that Vsp1-C301S injection did not affect the neural crest response towards SDF-1 coated beads (**Supplementary Fig.. 6; Supplementary Video 8**). Together, these data sets indicate that, while not being required for neural crest motility or chemotaxis, Vsp1 is essential for the response of the neural crest to electric fields and in turn for collective electrotaxis *in vivo*. Since the role of Vsp1 in electrotaxis has not been previously described, we predict that its study can have major implications across research fields. Indeed, by accessing publicly available databases, we found that orthologous *VSP* is expressed in several mammalian cell types, which are used as wound healing and cancer models and that have been shown to electrotax at least *in vitro* (**Supplementary Table. 2**), suggesting a requirement of Vsp1 in these biological contexts.

Our next goal was to explore the mechanism underlying the emergence of the endogenous electric fields^33^. Previous work has shown that membrane stretching activates ion channels to allow for ion translocation^34^. In addition, stress measurements in amphibians detected an increase of anisotropic tension in the neural plate, but isotropic tension in the non-neural ectoderm^35–37^. Further work showed that this anisotropic tension arises in the neural plate owing to the activity of the planar cell polarity (PCP) pathway in promoting convergent extension movements of the ectoderm^38, 39^. Thus, we hypothesised that the outward currents recorded in the neural fold, and in turn electric fields, emerge from ionic movement due to the stretching of neural fold cell membranes, owing to PCP-mediated convergent extension movements within the ectoderm. To test this hypothesis, we first confirmed that there was a tension build-up in the membrane of neural fold cells from stages in which outward electric currents were minimal or undetected (stage 13) to stages in which we recorded high levels of outward electric currents (stage 17), using laser ablation experiments (**Fig. 3a,b; Supplementary Fig.. 8; Supplementary Video 9)**. As this increased tension correlated with the emergence of electric fields, we next analysed the impact of PCP inhibition in neural fold to membrane tension and electric field formation (**Fig. 3a**). PCP was inhibited by targeted injections of DshDEP^+^, a known PCP inhibitor^39^, into the neural fold (**Supplementary Fig.. 7**). DshDEP^+^ injection led to a significant decrease in neural fold membrane tension (**Fig. 3b; Supplementary Fig.. 8; Supplementary Video 9**), as well as to a drastic reduction on the neural fold outward currents that, in turn, affected the establishment of endogenous electric fields (**Fig. 3c,d; Supplementary Fig.. 8**). Thus, we next tested whether the activity of stretch-activated channels was required for the formation of electric fields. For this, we performed live-current recordings before, during and after GsMTx4 inoculation, a peptide toxin which specifically inhibits stretch-activated channels in *Xenopus* and in other species^40, 41^ (**Fig. 3e**). We observed that shortly after GsMTx4 inoculation, the outward currents significantly decreased, abolishing the establishment of endogenous electric fields when comparing to the control (**Fig. 3f**). Furthermore, we found that neural crest cells dCCM *in vivo* was also impacted by the inhibition of PCP in the neural fold (**Supplementary Fig.. 7**), alike to the phenotype observed upon Vsp1 loss-of-function (**Supplementary Fig.. 5**). Importantly, this impact in migration is non-cell autonomous, since wild-type neural crest cells fail to directionally migrate once grafted into a host expressing DshDEP^+^ in the neural fold (**Fig. 3g–j; Supplementary Video 10**). Combined, our experiments show that PCP-mediated neural fold cell membrane stretching is required to the emergence of electric fields in the embryo and, consequently, to the directed migration of neural crest cells.

**Fig. 3.**
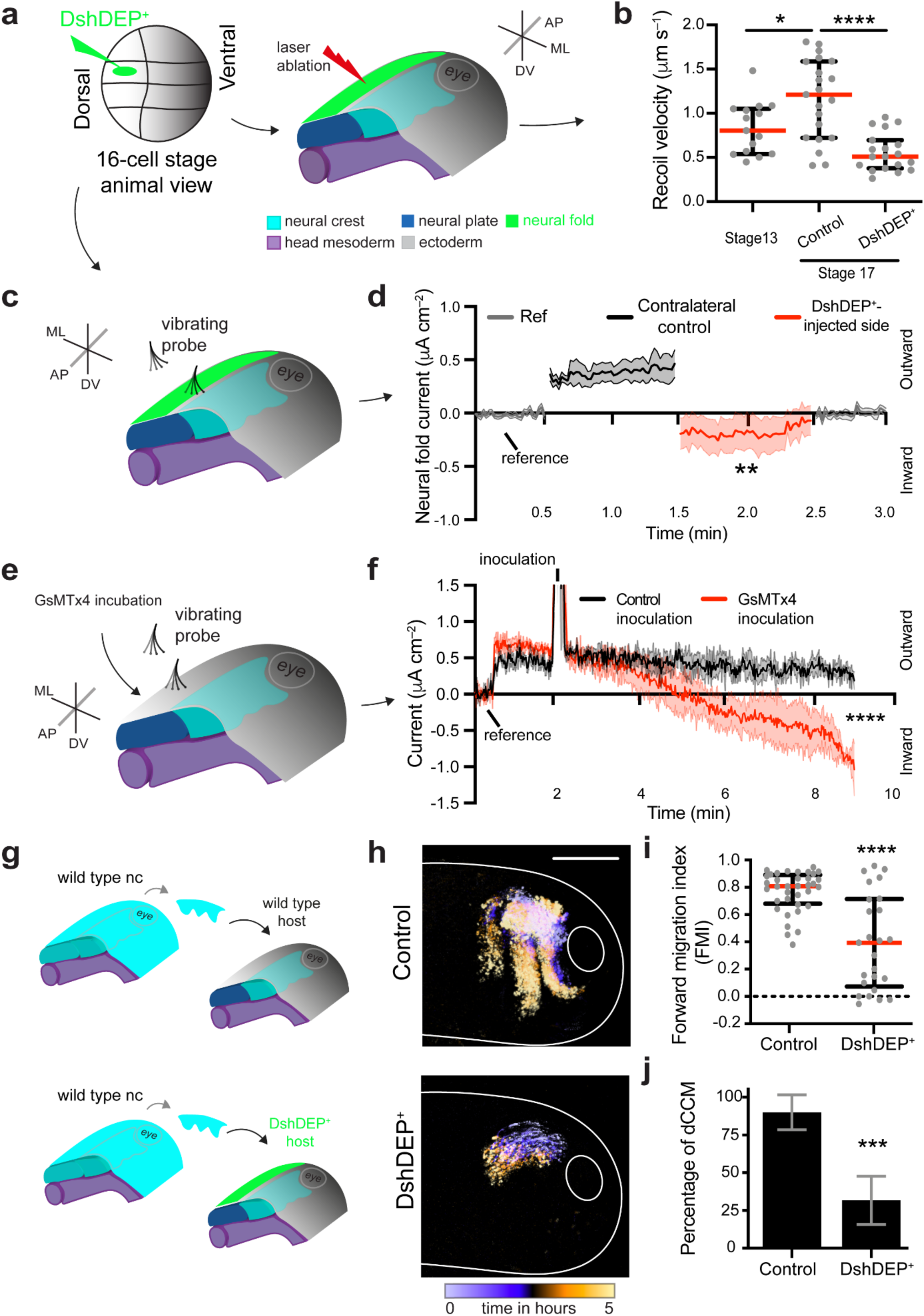
**Electric fields emerge from PCP-dependent neural fold membrane stretching to guide dCCM *in vivo*. a**, Scheme of DshDEP^+^ neural fold-targeted injection for subsequent laser ablation. (**b**) Neural fold membrane tension at non-migratory (st13) and pre-migratory (st17, control or DshDEP^+^ neural fold injected) stages. Red lines represent median and error bars the interquartile ranges, Student’s *t*-test, \**p* = 0.0180 (two-tailed), *t*-test with Welch’s correction, \*\*\*\**p* < 0.0001 (two-tailed); *n*_Non-migratory_ = 15 cell membranes, *n*_Control_ = 21 cell membranes, *n*_DshDEP_^+^ = 19 cell membranes. **c**, Scheme of vibrating probe measurement in DshDEP^+^ neural fold injected embryos. **d**, Impact of DshDEP^+^ neural fold targeted injections in electric currents, solid lines represent mean and shades the standard error, paired *t*-test (matched means of individual scatter lines), \*\**p* = 0.0019 (two-tailed), *n* = 14 embryos. **e**, Scheme showing the simultaneous GsMTx4 inoculation and electric currents measurements. **f**, Impact of Control or GsMTx4 (5 µM) inoculation during live recording of currents in the neural fold. (reference, probe away from embryo). Solid line represents mean and shades the standard error, Wilcoxon matched-pairs signed rank test (time-matched after inoculation), \*\*\*\**p* < 0.0001, *n* = 6 embryos. **g–j**, PCP inhibition in the neural fold impairs neural crest directionality. **g**, Scheme of the transplantation assays (nc, neural crest). **h**, Time colour-coded trajectories of neural crest cells (lateral view of embryos) (see **Supplementary Video 10**). Scale bar, 200 μm. **i**, Forward migration index (FMI) quantifications, red lines represent median and error bars the interquartile ranges, Mann Whitney *U*-test, \*\*\*\**p* < 0.0001 (two-tailed), *n*_Control_ = 36 cells, *n*_DshDEP_^+^ = 25 cells. **j**, Percentage of embryos displaying streams, column bars represent mean and error bars the standard deviation, Fisher’s exact test, \*\*\**p* = 0.0002 (two-tailed), *n*_Control_ = 19 embryos, *n*_DshDEP_^+^ = 18 embryos. **g**, Representative examples from at least three independent experiments; CI = 95%.

Finally, to confirm that the phenotype in the neural crest dCCM is a consequence of the effect of PCP in electric field formation, we assessed whether the introduction of a parallel exogenous electric field of identical magnitude and direction to the ones recorded *in vivo* would rescue PCP associated defects (**Fig. 4a,c**). In the absence of a parallel electric field, PCP inhibition in the neural fold non-autonomously inhibited neural crest dCCM *in vivo* (**Fig. 4a,b,e,f**), as we previously observed (**Fig. 3f,g**). Remarkably, the introduction of the parallel electric field rescued the dCCM of wild-type neural crests in these embryos (**Fig. 4c–f**). This electric field-induced rescue was lost when Vsp1-C301S was injected in the neural crest, further confirming the requirement of Vsp1 in the response to electric fields (**Fig. 4c–f**). These results confirm the role of the recorded electric fields in guiding neural crest dCCM via electrotaxis *in vivo* and the requirement of Vsp1 as an important part of the electrosensitive cascade by which neural crest cells transduce these endogenous electric fields into dCCM.

**Fig. 4.**
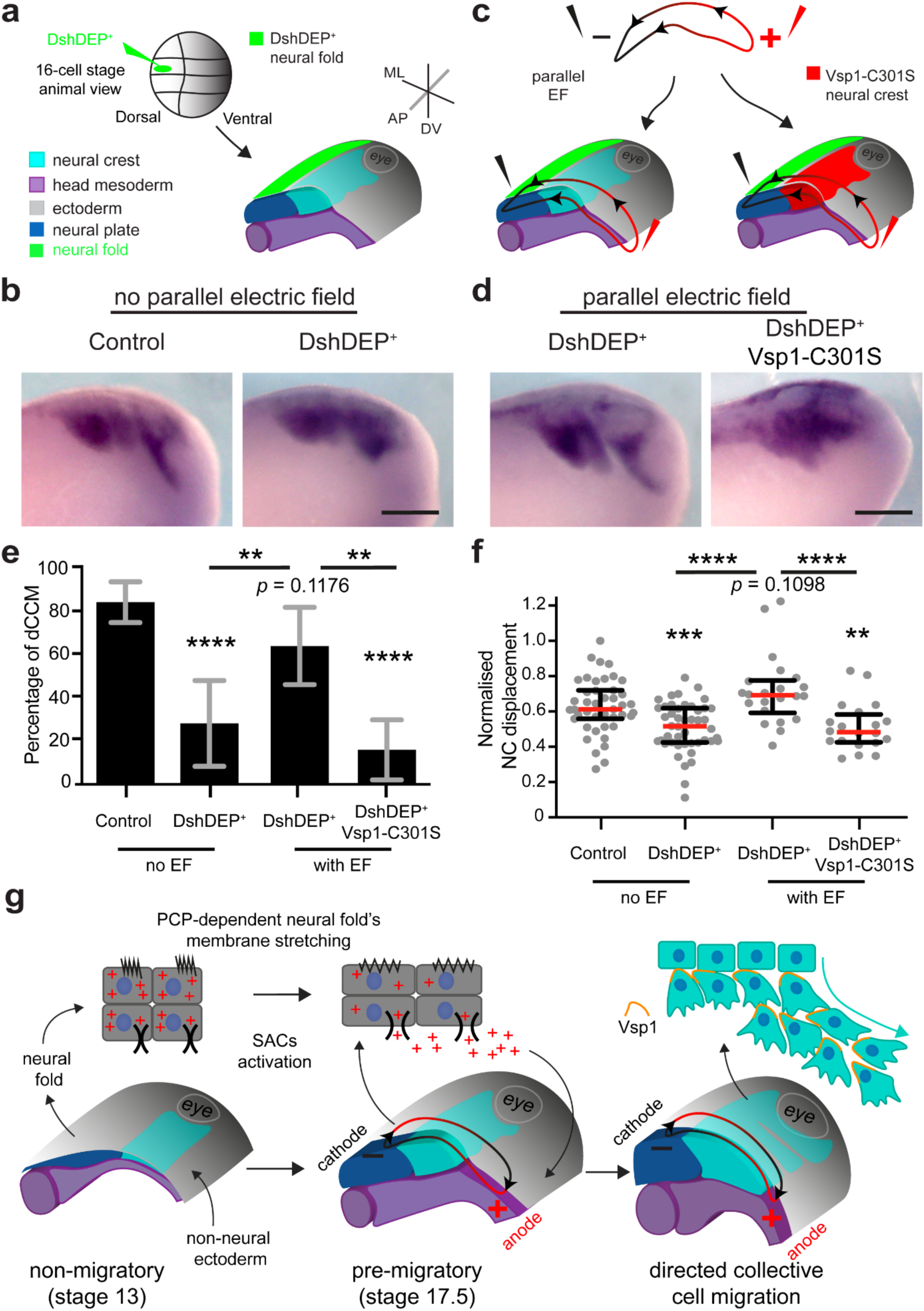
**Application of parallel electric fields rescues PCP-dependent impaired collective cell migration *in vivo*. a**, Scheme of DshDEP^+^ targeted injection into the neural fold. **b,** *In situ* hybridisations showing the effect of this injection in embryos in which no parallel electric fields (EFs) have been applied (lateral view of embryos hybridised with *sox*8, a neural crest migration marker). Scale bar, 200 μm. **c**, Scheme of the application of parallel EFs (100 mV mm^−1^) in embryos with DshDEP^+^ injected neural fold or in embryos with both DshDEP^+^-injected neural folds and Vsp1-C301S-injected neural crests. **d**, *In situ* hybridisations showing the effect of these treatments. Scale bar, 200 μm. **e**, Percentage of embryos displaying streams, column bars represent mean and error bars the standard deviation, Fisher’s exact test (two-tailed): \*\*\*\**p* < 0.0001 for Control vs. DshDEP^+^; *p* = 0.1176 for Control vs. DshDEP^+^ + EFs; \*\**p* = 0.008 for DshDEP^+^ vs. DshDEP^+^+ EFs; \*\*\*\**p* < 0.0001 for Control vs. DshDEP^+^ + Vsp1-C301S + EFs; \*\**p* = 0.0058 for DshDEP^+^ + EFs vs. DshDEP^+^ + Vsp1-C301S + EFs. **f**, Neural crest streams normalised displacement, red lines represent median and error bars the interquartile ranges, Mann Whitney *U*-test, (two-tailed): \*\*\**p* = 0.0004 for Control vs. DshDEP^+^; *p* = 0.1098 for Control vs. DshDEP^+^ + EFs; \*\*\*\**p* < 0.0001 for DshDEP^+^ vs. DshDEP^+^+ EFs; \*\**p* = 0.0023 for Control vs. DshDEP^+^ + Vsp1-C301S + EFs; \*\*\**p* = 0.0001 for DshDEP^+^+ EFs vs. DshDEP^+^ + Vsp1-C301S + EFs; *n*_Control_ = 43 embryos, *n*_DshDEP_^+^ = 46 embryos, *n*_DshDEP_^+^ _+ EFs_ = 22 embryos, *n*_DshDEP_^+^ _+ Vsp1-C301S + EFs_ = 20 embryos. **b,d**, Representative examples from at least three independent experiments; CI = 95%. **g**, Schematic summarising the mechanism of neural crest collective electrotaxis discovered here: PCP-driven membrane stretch allow for ion mobilization in the neural fold. This, in addition to physiological inward currents in the flanking non-neural ectoderm, allow the establishment of a sustained electric field along the migratory path of neural crest cells (cyan). Then, the activity of Vsp1 is required in the neural crest to specifically transduce these electric stimuli into dCCM.

A large body of *in vitro* evidence supports the idea that electrotaxis directs cell migration^17, 24^. *In vivo*, some hypotheses have been proposed to explain the establishment of endogenous electric fields^33^, although no experimental demonstration was provided to date. Here, we unveiled that endogenous electric fields are established in the migratory path of neural crest cells by PCP-driven mechanical stretching of the neural fold cell membranes. Furthermore, we demonstrated that the specific activity of Vsp1 in neural crest cells converts these endogenous electric fields into collective electrotaxis, which agrees with the topology of the electric fields recorded in the migratory path of neural crest cells *in vivo* (**Fig. 4g**). Thus, our research contributes to advance the accelerating field of electrotransduction^24, 25, 42^ by positioning Vsp1 as an electrosensor/transducer that has the potential to operate in several biological contexts (**Supplementary Table. 2**). More broadly, our work emphasises the idea that the occurrence of morphogenetic events such as dCCM are synchronised by the interplay between surrounding tissues, not only at the molecular, but also at the electrical and mechanical levels.

Thus, given the wide relevance of bioelectricity and directed cell migration in a variety of biological processes such as embryogenesis, tissue repair and cancer^1, 6, 16, 43, 44^, we believe that our results can deeply impact the research across these fields. Moreover, our research has the potential to influence the growing field of bioartificial organ production^45^, as this field may benefit from the inclusion of endogenous electric fields and from considering mechanoelectrical interplay in their protocols.

## Acknowledgments

The authors would like to thank Dr Benjamin Steventon (Cambridge University), Dr Ana Patrícia Ramos (IGC) and Dr Eric Theveneau (CBI-Toulouse) for helpful comments to the manuscript. In addition, the authors thank all lab members for helpful discussion, João Mata for technical assistance and Cristian Marchant for RNA extraction; Eric Theveneau for plasmids and *in situ* hybridization probes; and the IGC’s Advanced imaging (supported by PPBI-POCI-01-0145-FEDER-022122), Genomics, Bioinformatics and Aquatic animal facilities. We also thank Mr. Alan Shipley (Applicable Electronics, LLC.) for lending and assisting with the SVET system, and to Mr. Eric Karplus (Science Wares, Inc.) for software support. Work at E.H.B. lab was funded by an European Research Council (ERC) under the European Union’s Horizon 2020 research and innovation programme (grant agreement No. 950254), an EMBO IG Project Number 4765 and a la Caixa Junior Leader Incoming (94978). E.H.B. also acknowledge the support of Instituto Gulbenkian de Ciência and Fundação Calouste Gulbenkian (start-up fund I-411133.01). F.F. was supported by IGC (start-up fund I-411133.01) and an EMBO postdoctoral fellowship (ALTF 27-2020); S.M. was supported by an ERC-StG to E.H.B. and a FCT postdoctoral fellowship (2020.00759.CEECIND).

## Author contributions

F.F. and E.H.B. conceived the project and designed the experiments. F.F. performed most of the experiments and data analyses with help from E.H.B. S.M. performed RNA extractions, RNA-seq, PCR experiments and wrote methods for this part. F.F. and E.H.B wrote the manuscript and prepared all the figures. All the authors edited the manuscript.

## Competing financial interests

The authors declare non-financial competing interests.

## Supplementary information

Supplementary information is included here and videos are available upon request to the corresponding author.

## Methods

### Frog manipulation and embryo generation

*Xenopus laevis* (Daudin, 1802) embryos were obtained from *in vitro* fertilization as previously described^46^. Briefly, injection of human chorionic gonadotropin (MSD Animal Health, Chorulon) into the dorsal lymph sac was used to induce the ovulation in mature females. Upon gentle squeezing, the females laid oocytes that were fertilized using a sperm solution containing 1/5 of freshly collected testis in 500 µl of Marc’s modified Ringer 0.1× medium (MMR: NaCl 10 mM, CaCl_2_·2H_2_O 0.2 mM, KCl 0.2 mM, MgCl_2_·6H_2_O 0.1 mM, and HEPES 0.5 mM, adjusted to pH 7.1–7.2). To collect testes, mature males were dissected after being euthanized in overdose tricaine. Embryos were incubated between 12 and 23 °C until the desired stages were reached. *Xenopus* embryos were staged by following established developmental tables^47^. All animal procedures and euthanasia were reviewed and approved by the Ethics Committee and Animal Welfare Body (ORBEA) of the Instituto Gulbenkian de Ciência (IGC) and complied with the Portuguese (Decreto-Lei n° 113/2013) and European (Directive 2010/63/EU) legislation.

### Microinjection and pharmacologic modulations

Morpholino, mRNA and/or DNA injections were performed by targeted microinjections using calibrated glass needles that were mounted onto a cell microinjector (MDI, PM1000). Eggs were de-jellied in a cysteine solution (1 g of cysteine and 500 µl of NaOH 5 M in 50 ml MMR 0.1×) and transferred to ficoll 5% in MMR 0.45× (w/v) (Sigma-Aldrich, P7798). To target the neural crest population, one dorsal and one ventral animal blastomeres were injected at 8-cell stage (this injection can also target other ectodermal tissues). The neural fold was targeted by injecting one dorsal animal blastomeres at 16-cell stage (**Supplementary Fig.. 7**). Blastomeres were injected with 10 nl of the specified solution. When required, cells were fluorescently tagged with mRNA of nuclear RFP, membrane RFP and/or membrane GFP; transcripts were generated *in vitro* by using the mMESSAGE mMACHINE SP6 kit (Thermo-Fisher, AM1340), according to manufacturer’s instructions; and, 250 pg per embryo per construct was injected or co-injected with other treatments. The PCP inhibition was achieved with the injection of 2 ng DshDEP^+^ per embryo. The plasmids for Vsp1-C301S (Addgene, 51882), Vsp1-R152Q-GFP (Addgene, 51884), Vsp1-GFP (Addgene, 51883) and ElectricPk-cpEGFP (Addgene, 40314) were previously reported^29, 48^. Vsp1-C301S construct was injected for a final 1.5 ng per embryo. Vsp1-R152Q-GFP, Vsp1-GFP and ElectricPk-cpEGFP plasmids were transcribed using the mMESSAGE mMACHINE T7 kit (Thermo-Fisher, AM1344) and mRNA was injected for a final 1, 1.5 or 1.5 ng per embryo, respectively.

For drug modulation, the peptide toxin GsMTx4 (Smartox Biotechnology, 08GSM001) was dissolved in H_2_O in a stock of 200 µM and stored at −80 °C. GsMTx4 was reconstituted in MMR 0.1× and inoculated at a final concentration of 5 µM. The control of GsMTx4 was the inoculation of H_2_O diluted in MMR 0.1×.

### Neural crest dissection, culture and transplantations

Cephalic neural crest populations were dissected from embryos at stages 16 and 17 and cultured *ex vivo* as previously described^49^. Briefly, the vitelline membrane was removed with fine forceps (Dumont, #4) and the embryos were gently immobilized in modelling clay. A hair knife (eyebrow glued into a glass pipette) was used to remove the epidermis and carefully explant the underlying neural crest. These explants were cut into small clusters and cultured in Danilchik’s for Amy 1× medium (DFA: NaCl 53 mM, Na_2_CO_3_ 5 mM, potassium gluconate 4.5 mM, sodium gluconate 32 mM, MgSO_4_·7H_2_O 1 mM, CaCl_2_·2H_2_O 1 mM, BSA 0.1% (w/v), adjusted to pH 8.3 with Bicine 1 M). Clusters were then plated into glass-bottom dishes coated with 62.5 μg ml^−1^ of fibronectin (Sigma-Aldrich, F1141) and let to adhere to the substrate for 15–30 min. Typically, cultures were made from neural crests explanted from 3–6 embryos per condition and arranged in quincunxes.

For *in vivo* transplantation, appropriate wild-type or injected neural crests from donor and host were explanted as aforementioned. Using the hair knife, the donor neural crest was grafted into the host embryo and held in place by a small piece of coverglass (∼1×1 mm). Once host embryos healed (15–60 min), the glass was removed and embryos were cultured in MMR 0.3× until the desired stage. For time-lapse imaging, embryos were mounted on agarose dishes with ∼1 mm lanes filled with methyl cellulose 3% in phosphate-buffered saline (PBS; w/v) to minimize embryo drifting.

### Vibrating probe measurements *in vivo*

Extracellular net electric current density was measured using a non-invasive vibrating voltage probe as previously described^50, 51^. The vector form of Ohm’s law, *EF = J_I_ × ρ*, defines current density (*J_I_*) as directly proportional to electric fields (*EF*), with the resistivity (*ρ*) of the medium as the constant of proportionality^17^. Hence, electric currents and fields are intrinsically interdependent vector phenomena, with both magnitude and direction.

The measuring probe is a platinum/iridium microelectrode (MicroProbes for Life Science, PI10036.0A10) that was incorporated into the turnkey system Scanning Vibrating Electrode Technique (SVET; Applicable Electronics). Prior to measurements, the probe was platinum-electroplated at the tip (∼25 μm diameter) and calibrated with a known electric current density. Briefly, with the probe horizontally vibrating between 100–200 Hz, a glass microelectrode point source positioned 150 μm away and passing 60 nA calibrated the vibrating probe in MMR 0.1× for an electric current density of 21.2 μA cm^−2^.

The SVET system is fitted with a stereoscope that allows to position embryos into a customized non-conductive measuring chamber, containing a plastic lane to hold the embryos in a dorsal up position, allowing perpendicular access of the vibrating probe. The currents were measured 10–30 μm away from the embryo surface and sampled every 1 s until a plateau was reached (typically in <1 min) (**Supplementary Fig.. 1**). Reference values were acquired with the probe >1 mm away from embryo. As we follow the conventional current, defined by the flow of positive charge, outward current density is the net positive charge exiting the embryo and inward current density is the one entering the embryo. Measurements in the regions and stages specified occurred at 20–22 °C in MMR 0.1×. Data and metadata were acquired and extracted using the interface software Automated Scanning Electrode Technique (ASET-LV4; Science Wares) and statistically analysed using Excel 2016 (Microsoft) and GraphPad Prism v9.0.0 (GraphPad Software).

### Electric field strength estimation

The imposed EFs magnitude was estimated from the embryo currents that we measured during neurulation. We observed differential current drops (neural fold minus flank non-neural ectoderm) ranging ∼0.5–2.5 µA cm^−2^, over an approximate 200–400 µm distance (**Fig. 1b–d**). Following Ohm’s law, these currents should generate ∼4–12 mV voltage drops, corresponding to mediolateral endogenous EFs of ∼10–60 mV mm^−1^ detected along the migratory path of neural crest cells. These values fall within previously determined EFs of ∼5–75 mV mm^−1^ in rostral regions of different amphibians^13, 14^. Thus, we followed these values in our *ex vivo* and *in vivo* electrotaxis assays.

### Electrotaxis assay *ex vivo*

The application of exogenous direct current electric fields^52^ was performed in a custom-made electrotaxis chamber. The electrotactic chambers with a 22×8×0.3 mm EFs tunnel were created by placing two single-use strips of polydimethylsiloxane (PDMS) silicone (Sylgard, 184) (0.3 mm height) onto the sides of a reusable 1-well chambered coverglass (Nunc, 155360). For every new assay, the bottom coverglass was replaced by an equivalent coverglass glued with twinsil silicone (Picodent, 1300 1000). To build the PDMS strips, PDMS was prepared by mixing the elastomer/curing agent in a 10:1 ratio. This mix was vacuum degassed and poured into a mould assembled from two silanized microscopic slides spaced by two cover glasses no. 1. The PDMS was cured overnight at 65 °C, manually trimmed and cut into strips. At this stage, neural crest clusters were plated into the fibronectin-coated electrotactic chamber tunnel and let to adhere (**Fig. 1e**). Then, using electric insulating grease (Dow Corning, DC4), a coverglass roof was attached to the PDMS sides to cover the clusters and electrically seal the EF tunnel. Two dams made of PDMS (18×2×7 mm) were sealed with grease to the extremes of the roof, generating DFA 1× medium reservoirs with 3 ml volume capacity. To minimize evaporation, a lid with holes for the agar salt bridges and voltmeter access was used. This setup was mounted for time-lapse microscopy and once all positions were selected, EFs of the above calculated magnitudes were applied. The EFs were channelled through the tunnel via chloritized silver electrodes (Advent, AG549107) and agar salt bridges, composed of agarose 1.5% in Steinberg’s solution (w/v) (NaCl 58 mM, KCl 0.67 mM, MgSO_4_·7H_2_O 1.3 mM, CaNO_3_·4H_2_O 0.44 mM, Tris base 4.6 mM, adjusted to pH 7.4). The required voltages were provided by a direct current power supply (Apelex, 117240 or BK Precision, 9132B) fine-tuned by a resistance decade box (Tenma, 72-7270) and measured in the electrotactic chamber by a voltmeter. Following conventional current, charge flows from anode (+ pole) to cathode (− pole).

### Application of electric fields *in vivo*

To apply parallel or antiparallel exogenous electric fields (100 mV mm^−1^) along the migratory path of neural crest cells, we used the same power supply, electrodes and agar bridges described in the *ex vivo* setup, but used a different electric chamber. Briefly, in a Petri dish half filled with modelling clay, a 22×10×1 mm EFs tunnel was modelled into the centre of the dish. Under a stereoscope, the embryos were gently immobilized in the clay, arranged in quincunxes and oriented to maximize the effect of the applied EFs. Coverglass roof, dams and lid were used as in the *ex vivo* electrotaxis assay. EFs were applied from stage 17 to stage 22 and embryos were immediately fixed and processed for *in situ* hybridization (as described below).

### Motility, protrusion and chemotaxis assays

To test the basal motility and directionality of wild-type and transfected neural crests, we used motility, protrusion and chemotaxis assays^43, 49^. For motility, the neural crest clusters were plated into a plastic-bottom dish (Falcon, 351006) coated with fibronectin 125 μg ml^−1^. After adhesion for 1 h, the tight-fit lid was carefully put and the dish was flipped for imaging. For protrusion, the clusters were plated into a fibronectin-coated glass-bottom dish and let to adhere prior to time-lapse microscopy. For chemotaxis, heparin-acrylic beads (Sigma-Aldrich, H5263 (discontinued)) were soaked overnight with human SDF-1 3.3 μg ml^−1^ (Sigma-Aldrich, S1577) at 4 °C. Using a fibronectin-coated glass-bottom dish (WPI, FD5040), the clusters were placed 2–3 diameters away from the beads that were fixed into a line of grease (**Supplementary Fig.. 6**).

### *In situ* hybridization

Embryos were fixed, dehydrated and rehydrated, bleached, hybridized, blocked and stained following *in situ* hybridization standard protocols^49^. Digoxigenin-labelled antisense probes against *complement component* 3 (*c*3)^53^ or *sox*8^54^, were used to recognise the neural crest cells in whole embryos. Probes were transcribed with a Riboprobe *in vitro* Transcription System (Promega, P1420), according to manufacturer’s instructions. The signal was revealed using the alkaline phosphatase detection reagents NBT/BCIP (Roche, 11383221001).

### *In situ* hybridisation imaging

*In situ* hybridized embryos were mounted on an agarose dish with small depressions to facilitate acquisition. An USB Dino-Eye eyepiece camera (Dino-Lite, AM7025X), mounted onto a stereoscope with cold light, was used to take photomicrographs at 2.5× magnification, using DinoCapture v2.0 (Dino-Lite).

### sqRT-PCR

Neural crest explants from pre-migratory stages (17–18) were collected and processed for RNA extraction using the RNeasy Mini kit (Qiagen, 74104). Briefly, 5 ng of isolated total RNA was used as a template for the RT performed using SuperScript IV (Invitrogen, 18090050), according to manufacturer’s instructions. To run the PCRs, 2.5 µl of resultant cDNA in 25 µl of reaction was used for the genes of interest, using Q5 Hot Start High-Fidelity DNA Polymerase (NEB, 174M0493S) with the following primers: *vsp*1 (F–GGAGTAATGAGTGCGACCGA; GGCTACGATGACTGTGCCAT; R–TCGACAGTAACAACTTGGTGA); *vsp*2 (F–TCATTGCTTTGTCTCCCTCCC; R–TGACTGAACCCTTCCACGA); *ef*1*α* (F–ACCCTCCTCTTGGTCGTTTT; R–TTGGTTTTCGCTGCTTTCT); *krt*12 (F–CACCAGAACACAGAGTAC; R–CAACCTTCCCATCAACCA); *sox*8 (F–GAATCATTTGCGCATTTGTGCC; R–AGTCACTTTGCCCTGCCTT); *xbra* (F–GGAGTAATGAGTGCGACCGA; R–GCCCGACATGCTCACCTTCA). PCR conditions were as follows: (1) 98 °C for 30 s (initial denaturation); (2) 98 °C for 10 s (denaturation); (3) 30 s (annealing) at 64 °C (*ef*1*α*), 60 °C (*krt*12), 66 °C (*sox*8), 64 °C (*vsp*1), 68 °C (*vsp*2) or 68 °C (*xbra*); (4) 72 °C for 20 s (34× cycles from step 2 to 4); and, finally, 72 °C for 2 min for final extension. The amplified fragments were resolved in agarose 0.8% (w/v) gel electrophoresis together with 1 Kb Plus Ladder (Invitrogen, 10787026). While *vsp*1, *vsp*2, *xbra* and *sox*8 primers were originally designed, *ef*1*α* and *krt*12 primers were previously published^55^.

### RNA-seq procedures and analysis

After assessing the quality of the extracted RNA using HS RNA Screen Tape Analysis (Agilent Technologies), mRNA-library was prepared using an adaptation of the SMART-Seq2^56^. Illumina libraries were performed using an adaptation of the Nextera protocol^57^. The library quantification and quality verification were done using the Fragment Analyzer (AATI) in combination with HS NGS Kit (Agilent Technologies). Sequencing was carried out in NextSeq500 Sequencer (Illumina) using 75 SE high throughput kit. Sequence information was extracted in FastQ format using the bcl2fastq v2.19.1.403 (Illumina).

The RNA-seq analysis was a service conducted by the Bioinformatics Unit at IGC. Briefly, reads from 18S and 28S rRNA were identified and discarded. The good-quality non-ribosomal reads (50585 in 51061) were mapped against the reference genome of *Xenopus laevis* using the annotation XENLA_9.2_Xenbase.gtf (v9.2). The genome FASTA and GTF files were downloaded from Xenbase. The gene expression tables were imported into the R v3.6.3 (The R Foundation for Statistical Computing) to normalize gene expression with the TMM (Trimmed Mean of M-values) procedure^58, 59^ with the NOISeq R package (v2.30.0)^60^.

### Localization and activation of Vsp1

To assess the cellular localization and activity of Vsp1, we injected Vsp1-GFP and ElectricPk-cpEGFP, respectively. For localization, neural crest clusters were injected with Vsp1-GFP as described above and plated into fibronectin-coated glass-bottom dishes (WPI, FD35) and let to adhere prior to confocal microscopy. To report Vsp1 response to electric fields we used the Vsp1-based sensor ElectricPk-cpEGF. This sensor was co-injected with the membrane RFP as described above and the expressing clusters without application of EFs or exposed to EFs of 100 mV mm^−1^ for 1 h were fixed overnight in formaldehyde 3.7% in PBS (v/v). After fixation, clusters were washed with PBS and mounted with mowiol (Millipore, 475904) and sealed with nail polish for confocal microscopy.

### Cryosectioning

Embryo sectioning was performed as previously described^43^. Briefly, neural fold-injected embryos were fixed for 2 h at room temperature and washed twice with phosphate buffer 1× (NaH_2_PO_4_·H_2_O 0.2 M and K_2_HPO_4_ 0.2 M, pH 7.4) for 5 min each wash. Embryos were passed to sucrose 15% in phosphate buffer (w/v) for 2 h at room temperature. Embryos were then embedded and orientated in a gelatine solution (gelatine 8% in 15% sucrose in phosphate buffer, w/v) and incubated for 1 h at 42 °C. Gelatine blocks were flash frozen at −80 °C in pre-cooled heptane and sectioned in 25 μm slices in a cryostat (Leica, CM3050 S). The slides dried overnight at room temperature and were mounted with mowiol and sealed with nail polish prior to confocal microscopy.

### Laser ablation assay

Recoil velocity as a proxy of membrane tension build-up or stretching was assessed using laser ablation experiments^61^. The centre of fluorescently labelled and mediolaterally oriented membrane junctions (i.e., perpendicular to the neural fold) was photoablated using a MicroPoint system (Andor) fitted with a 350 nm pulsed laser and 365/435 dye. The system was controlled by Metamorph software (Molecular Devices). Junctions were ablated using one pulse of 5–10% laser at 21–22 °C after the first intact image was acquired. Up to 3 ablations were performed per embryo, with sufficient distance from each other to avoid the effect of local membrane relaxation.

### Time-lapse imaging and confocal laser scanning

#### Graft experiments

Grafted embryos were *z*-stacked (10 µm sections) and imaged using 488 and 561 nm laser lines on an Imager Z2/ApoTome.2 system (Zeiss) equipped with an Orca Flash 4.0 v2 CMOS camera (Hamamatsu), or on a SP5 confocal system (Leica) with spectral detection. Water immersion 10× objectives were used in both systems: N-Achroplan 10×/0.30 NA (Zeiss) and HC APO L 10×/0.30 NA (Leica), respectively. Microscopes (both upright setups) were controlled by ZEN v3.1 (Zeiss) or LAS AF (Leica), respectively. Recording started at around stage 20 and was acquired every 5 or 10 min for up to 7 h at 21–22 °C.

#### Electrotaxis

Cell migration of neural crest clusters was recorded on a Nikon HCS microscope, equipped with a Zyla 4.2 sCMOS camera (Andor) and using a PL APO 10×/0.45 NA objective (Nikon). The system was controlled by NIS-Elements (Nikon) to acquire time-lapse microscopy every 5 min for up to 10 h at 18 °C, using 488 and 561 nm filter sets.

#### Motility

The inverted dish was mounted in the ApoTome system and clusters were imaged using the EC Plan-Neofluar 10×/0.30 NA objective by time-lapse mode every 5 min for up to 8 h at 21–22 °C.

#### Protrusions

The clusters were imaged using the SP5 confocal system with the HC PL APO 40×/0.80 NA water immersion objective (Leica). Recording occurred in time-lapse mode every 30 or 60 s for up to 30 min at 21–22 °C.

#### Chemotaxis

Cell migration was imaged using the SP5 confocal system or a Thunder Imager 3D Cell Culture system (Leica) with the HC PL APO 10×/0.45 NA objective (Leica), in time-lapse mode every 5 min for up to 6 h at 18–22 °C.

#### Laser ablation

Embryos were imaged using a HC PL APO 63×/1.30 NA glycerine immersion objective (Leica) mounted in an inverted confocal microscope (Yokogawa CSU-X Spinning Disk confocal, onto a Leica DMi8 microscope body). The system was controlled by Metamorph to acquire time-lapse imaging every 2 or 5 s for up to 30 s, using 488 and 561 nm laser lines and an iXon Ultra EMCCD camera (Andor).

#### Vsp1-GFP imaging

Injected clusters were imaged using an inverted confocal system (Zeiss, LSM 980), equipped with two PMT and one GaAsP and controlled by ZEN Blue v3.0 (Zeiss). Using the 488 and 561 nm laser lines, a *z*-stack (1 µm sections) was acquired at 21–22 °C with the C Plan-Apochromat 63×/1.40 NA oil immersion objective (Zeiss).

#### ElectricPk-cpEGFP imaging

Injected clusters were imaged using an upright confocal system (Leica, Stellaris 5), equipped with Power HyD S detectors and controlled by LAS X (Leica). Using the 488 and 561 nm laser lines, a *z*-stack (1 µm sections) was acquired at 21–22 °C with the HC PL APO 63×/1.40 NA oil immersion objective (Leica).

#### Neural fold-targeted injection imaging

Embryos of targeted injections of DshDEP^+^ into the neural fold were imaged using the Stellaris 5 confocal system with the HC APO L 10×/0.30 NA water immersion objective (Leica). The sections of these embryos were imaged using the LSM 980 confocal system with the LD LCI Plan-Apochromat 25×/0.80 NA oil immersion objective (Zeiss).

### Cell motility analysis

#### Cluster and cell migration

The centre of *ex vivo* clusters and *in vivo* cells were tracked typically for 3–6 h (exact time in figure legends) using the Manual Tracking plug-in in Fiji (https://imagej.net/Fiji; v1.53k). The cluster/cell directedness and velocity were computed using the Chemotaxis and Migration Tool v2.0 (Ibidi). Directedness is represented by the forward migration index (FMI), which shows how efficiently cells migrate forwardly parallel to the EFs vector *ex vivo* or mediolateral axis *in vivo*. The frequency of angles of the clusters/cells in relation to the EFs vector was computed using Rozeta v2.0 (freeware developed by Jacek Pazera). The anode was set at 180°. The frequency (percentage) of anodal migration was determined by counting the clusters that experienced a biased displacement of the whole cluster diameter towards the anode. In control conditions under EFs, clusters typically displaced several diameters (**Supplementary Fig.. 2, Supplementary Video 1,2**).

#### Collective vs. single cell migration

Electrotaxis time-lapses, from independent experiments, where several single cells detached from the migrating collective (cluster) were selected for comparative tracking. The centre of the cluster or single cells were tracked for 1 h. Tracking and analysis were performed as aforementioned.

#### Single cell motility

The cluster was let to disperse to allow the appearance of many motile single cells. Single cells were tracked for 1 h (**Supplementary Fig.. 5**) and analysed as above.

#### Cell protrusions

Maximum protrusion area was measured in Fiji using the freehand selection tool (**Supplementary Fig.. 5**). Protrusion dynamics correspond to the time for a complete cycle of protrusion expansion and retraction. Maximum protrusion area was determined at the maximum protrusion expansion frame.

### Neural crest migration analysis

The frequency of *in vivo* dCCM was determined by counting the *in situ* hybridized embryos which formed stereotypical streams (a hallmark of dCCM^18, 19^). In control conditions, majority of embryos formed three well-defined streams (**Fig. 1a,k**). The displacement of neural crest cells was derived with the ratio between the stream length and dorsoventral length of the embryo. All ratios were normalized against the maximum length of the control stream. Lengths were obtained in Fiji using the measurement tool.

### Membrane ablation analysis

The right and left vertexes of fluorescently labelled membranes were tracked for up to 20 s post photoablation using the Manual Tracking plug-in in Fiji (**Supplementary Fig.. 8**). The recoil velocities were averaged using Excel and plotted with GraphPad Prism.

### Image and video treatment

Substacking, *z*-projections (maximum intensity), rotations, time colour-coded projections, and time-lapse videos were performed in Fiji. General treatment, including adjustment of contrast and brightness, resizing, pseudocolouring, addition of scales and overlay of text in images and videos were conducted in Fiji and/or Photoshop 2021 (Adobe). Some of the time-lapses, particularly from grafts, were stabilized using the StackReg registration plug-in in Fiji. In time colour-coded projections, the background with debris and dead cells was coloured in black in Fiji, for clarity purposes and without interfering with the sample itself.

### Statistical analysis

The experimenters were not blinded in data acquisition, treatment, or statistical analysis due to the nature of the experimental procedures and the clear embryonic and cellular phenotypes of some manipulations. The embryos were randomly allocated to the specified experimental conditions. Data exclusion criteria were in place to filter out unviable, overaged, misinjected and contaminated embryos and/or cell clusters. Data outliers were removed based on the biological context when flagged using the statistical Grubbs’ test and/or ROUT method computed using GraphPad Prism. Majority of datasets presented no statistical outliers. The appropriate inferential statistics were computed using GraphPad Prism and are annotated in the figure legends. For parametric tests, pre-tests were conducted for the assumptions of normality (Kolmogorov-Smirnov and/or Shapiro-Wilk tests) and equal variances (F test). Majority of datasets verified these assumptions. To address non-verified assumptions, we used the Welch’s correction or a non-parametric equivalent test. Data representation and sample size (*n*, biological replicates) are annotated in figures legends. At least three independent experiments were performed per physiological readout, using embryo batches from different females. Differences were significant when the two-tailed *p* value was ≤ 0.05 with the following levels of significance: *p* > 0.05, non-significant, \**p* ≤ 0.05, \*\**p* < 0.01, \*\*\**p* < 0.001, and \*\*\*\**p* < 0.0001. Data analysis and visualization were performed using Excel, ASET, Chemotaxis and Migration Tool, Rozeta, R, GraphPad Prism, Fiji, Photoshop and Illustrator 2021 (Adobe).

### Data and code availability

The data supporting the findings of this study are embedded in the main text and supplementary information. In addition, other relevant data are available and unrestricted from the corresponding author upon reasonable request.

---

**Supplementary Fig. 1.**
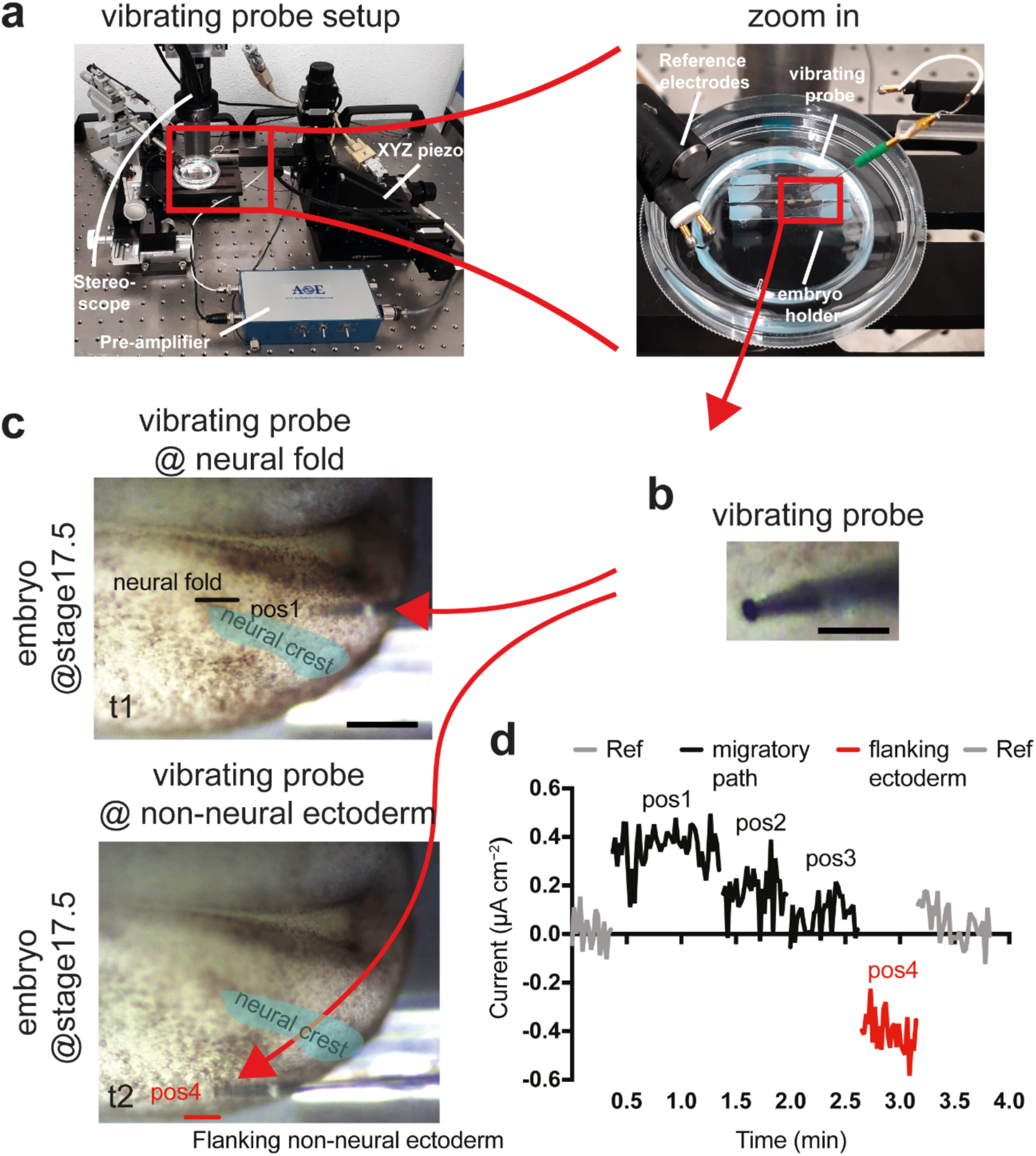
**Vibrating probe setup and measurements approach. a**, Overview of the vibrating probe setup to measure electric current densities *in vivo*. Zoom in of the custom-made measuring chamber. **b**, Detail of the probe in static mode, note the platinum black ball (∼25 µm diameter) electroplated to the tip. **c**, Representative photomicrographs acquired during a typical measurement; the probe in vibrating mode is indicated by a red arrow and shown in the neural fold (pos1) and then in the flanking non-neural ectoderm (pos4); other two records (pos2 and 3) were made between pos1 and 2, so the whole migratory path of neural crest cells was covered in each analysed embryo. Relative position of the neural crest is indicated in fading cyan. **d**, Representative result of the mediolateral records; pos1–pos4, positions along the migratory path, from neural fold to non-neural flanking ectoderm positions and Ref, reference (measurement obtained by holding the probe away from embryo).

**Supplementary Fig. 2.**
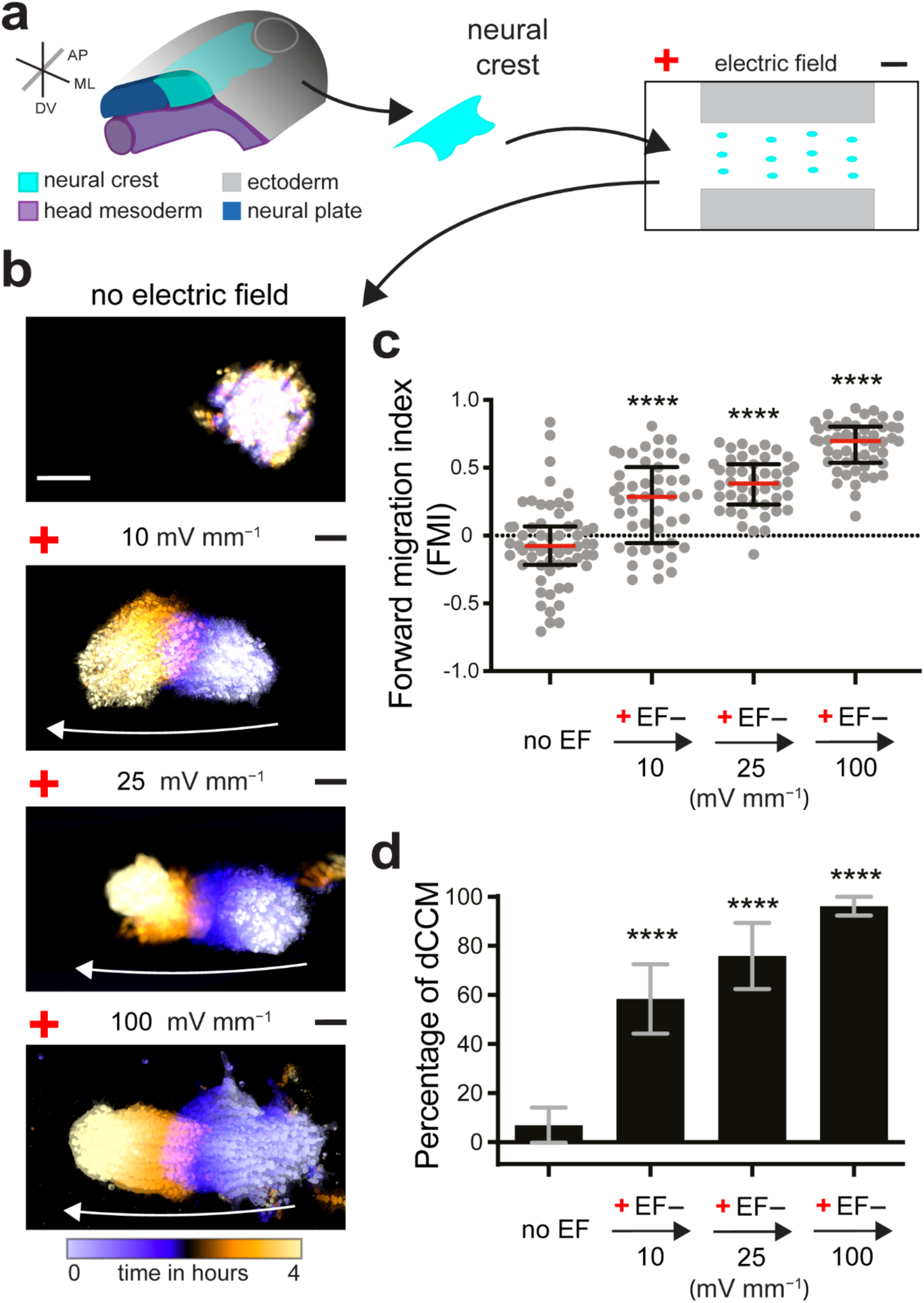
***Ex vivo* electrotactic response of neural crest cells at the different magnitudes of electric fields recorded *in vivo*. a**, Scheme of the electrotaxis assay. **b**, Time colour-coded trajectories of neural crest clusters, white arrows depict the cluster trajectories (see **Supplementary Video 2**). Scale bar, 100 μm. **c**, Forward migration index (FMI) quantifications, red lines represent median and error bars the interquartile ranges, One-way ANOVA with Dunnett’s multiple comparisons test, \*\*\*\**p* < 0.0001 (two-tailed) for all electric fields (EFs) vs. No EFs, *n*_No EFs_ = 59 clusters, *n*_10 mV mm_^−1^ = 47 clusters, *n*_25 mV mm_^−1^ = 44 clusters, *n*_100 mV mm_^−1^ = 50 clusters. **d**, Percentage of clusters displaying migration towards the anode; column bars represent mean and error bars the standard error, Fisher’s exact test, \*\*\*\**p* < 0.0001 (two-tailed) for all EFs vs. No EFs, *n*_No EFs_ = 59 clusters, *n*_10 mV mm_^−1^ = 47 clusters, *n*_25 mV mm_^−1^ = 45 clusters, *n*_100 mV mm_^−1^ = 51 clusters. **b,** Representative examples from at least three independent experiments; CI = 95%.

**Supplementary Fig. 3.**
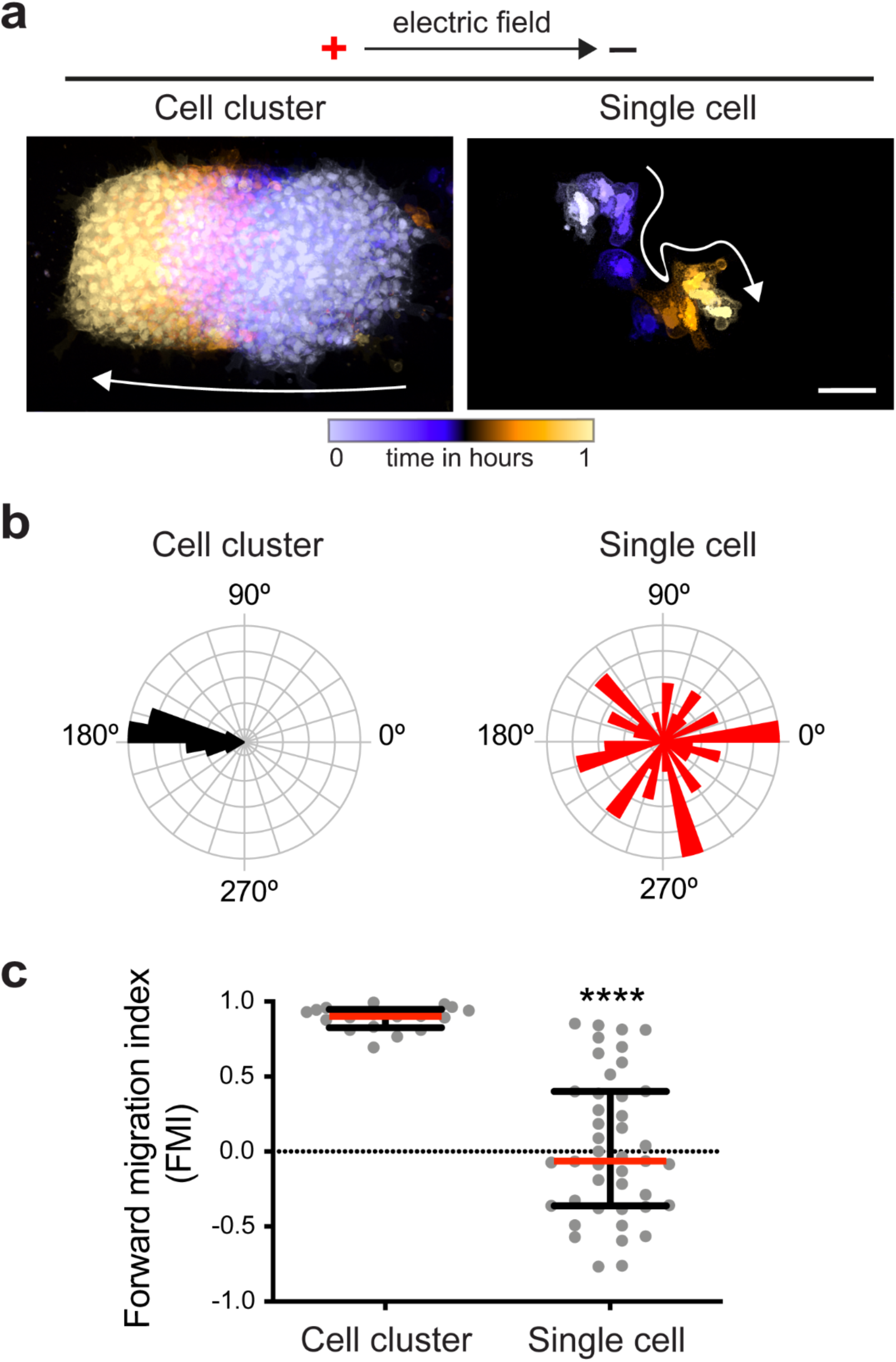
**Electrotaxis is an emergent property of cell clusters. a**, Time colour-coded trajectories of neural crest clusters and isolated neural crest cells migrating in electric fields of 100 mV mm^−1^, white arrows depict the cluster or cell trajectories (see **Supplementary Video 3**). Scale bar, 100 μm. **b**, Rose plots showing the angle frequencies of neural crest migration in relation to the electric field vector (anode at 180°). **c**, Forward migration index (FMI) quantifications, red lines represent median and error bars the interquartile ranges, Student’s *t*-test with Welch’s correction, \*\*\*\**p* < 0.0001 (two-tailed), *n*_Clusters_ = 18, *n*_Single cells_ = 43. **a,** Representative examples from at least three independent experiments; CI = 95%.

**Supplementary Fig. 4.**
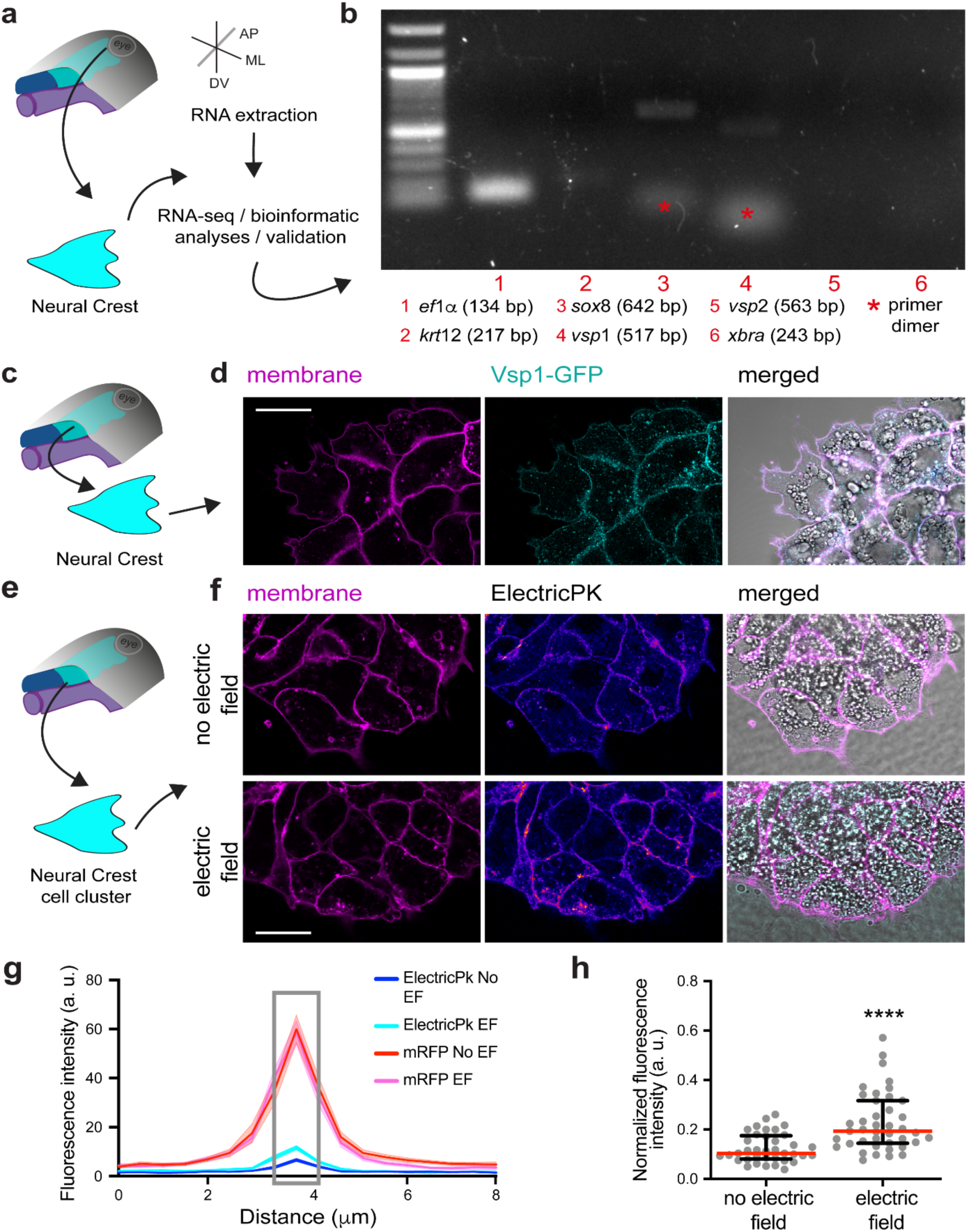
**Neural crest cells express the voltage-sensitive phosphatase, Vsp1. a**, Scheme and workflow of RNA extraction from neural crest explants dissected from pre-migratory embryos at st17–18 and processed for RNA-seq. **b**, *vsp*1, but not *vsp*2, is present in the neural crest cells. Purity of the explants further assessed by semiquantitative RT-PCR by confirming the presence of the neural crest marker (*sox*8) and the absence of ectodermal (*krt*12) and mesodermal (*xbra*) markers (ruling out contamination with surrounding tissues). **c**, Scheme of dissection of the neural crest. **d**, Membrane localization of Vsp1 in a neural crest cluster. **e**, Scheme of dissection of the neural crest. **f**, Activity of Vsp1 in neural crest clusters in the absence or presence of electric fields (EFs) of 100 mV mm^−1^ for 1 h as reported by the ElectricPk-cpEGFP construct. ElectricPk-cpEGFP intensity is colour-coded in fire lookup table (hot colours represent higher intensity). **g**, Fluorescence intensity profile of lines overlaying membranes, solid lines represent mean and shades the standard error. **h**, Maximum intensity normalized against the membrane RFP (grey rectangle in **g**). Red lines represent median and error bars the interquartile ranges, Mann Whitney *U*-test, \*\*\*\**p* < 0.0001 (two-tailed), *n*_No EF_ = 37 cell membranes, *n*_EF_ = 41 cell membranes. Scale bars in **d** and **f**, 20 μm. **b**,**d**,**f**, Representative examples from at least three independent experiments; CI = 95%.

**Supplementary Fig. 5.**
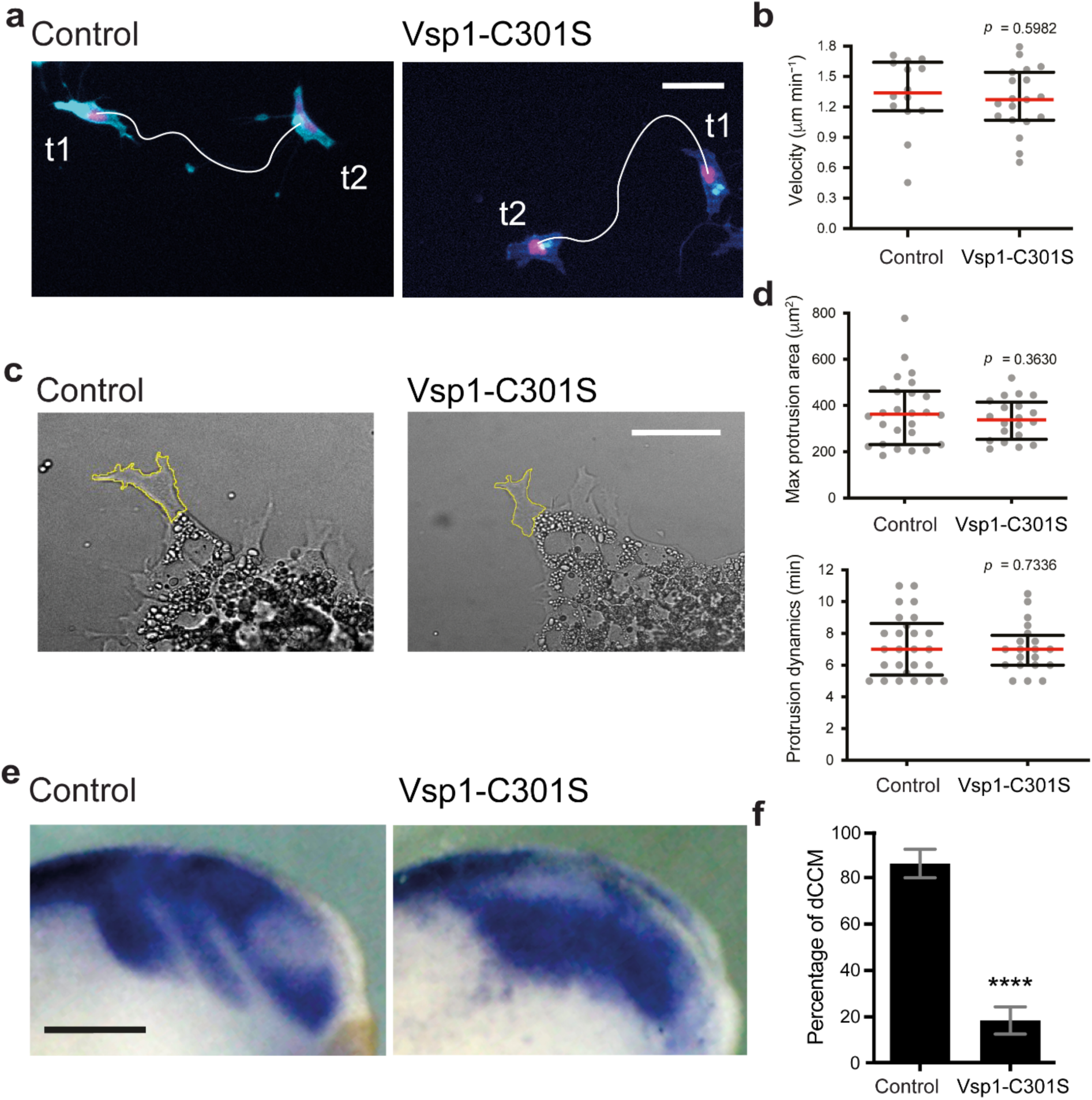
**Vsp1-C301S does not affect for neural crest cells motility *ex vivo* but it impairs stream formation *in vivo*. a**, Overlay of isolated neural crest cells at t1 (0 min) and t2 (60 min) of migration, solid white lines depict cell trajectories (see **Supplementary Video 5**). Scale bar, 30 μm. **b**, Quantification of single cells velocity, red lines represent median and error bars the interquartile ranges, *t*-test, *p* = 0.5982 (two-tailed), *n*_Control_ = 14 cells, *n*_Vsp1-C301S_ = 19 cells. **c**, Neural crest clusters with outlined protrusions (yellow solid lines). Scale bar 30 μm. Quantification of maximum protrusion area (**d**) and protrusion dynamics (**e**), red lines represent median and error bars the interquartile ranges, *t*-test (**d**) with Welch’s correction (**e**), *p*_Max Area_ = 0.3630 (two-tailed), *p*_Dynamics_ = 0.7336 (two-tailed), *n*_Control_ = 26 protrusions, *n*_Vsp1-C301S_ = 20 protrusions. **f,g**, Vsp1-C301S impairs stream formation (a hallmark of dCCM) *in vivo*. **f**, *In situ* hybridisations showing lateral views of embryos hybridised with *c*3 (a marker of neural crest migration). Scale bar, 200 μm. **g**, Percentage of embryos displaying stream formation, column bars represent mean and error bars the standard error, Fisher’s exact test, \*\*\*\**p* < 0.0001 (two-tailed), *n*_Control_ = 25 embryos, *n*_Vsp1-C301S_ = 24 embryos. **a**,**c**,**f**, Representative examples from at least three independent experiments; CI = 95%.

**Supplementary Fig. 6.**
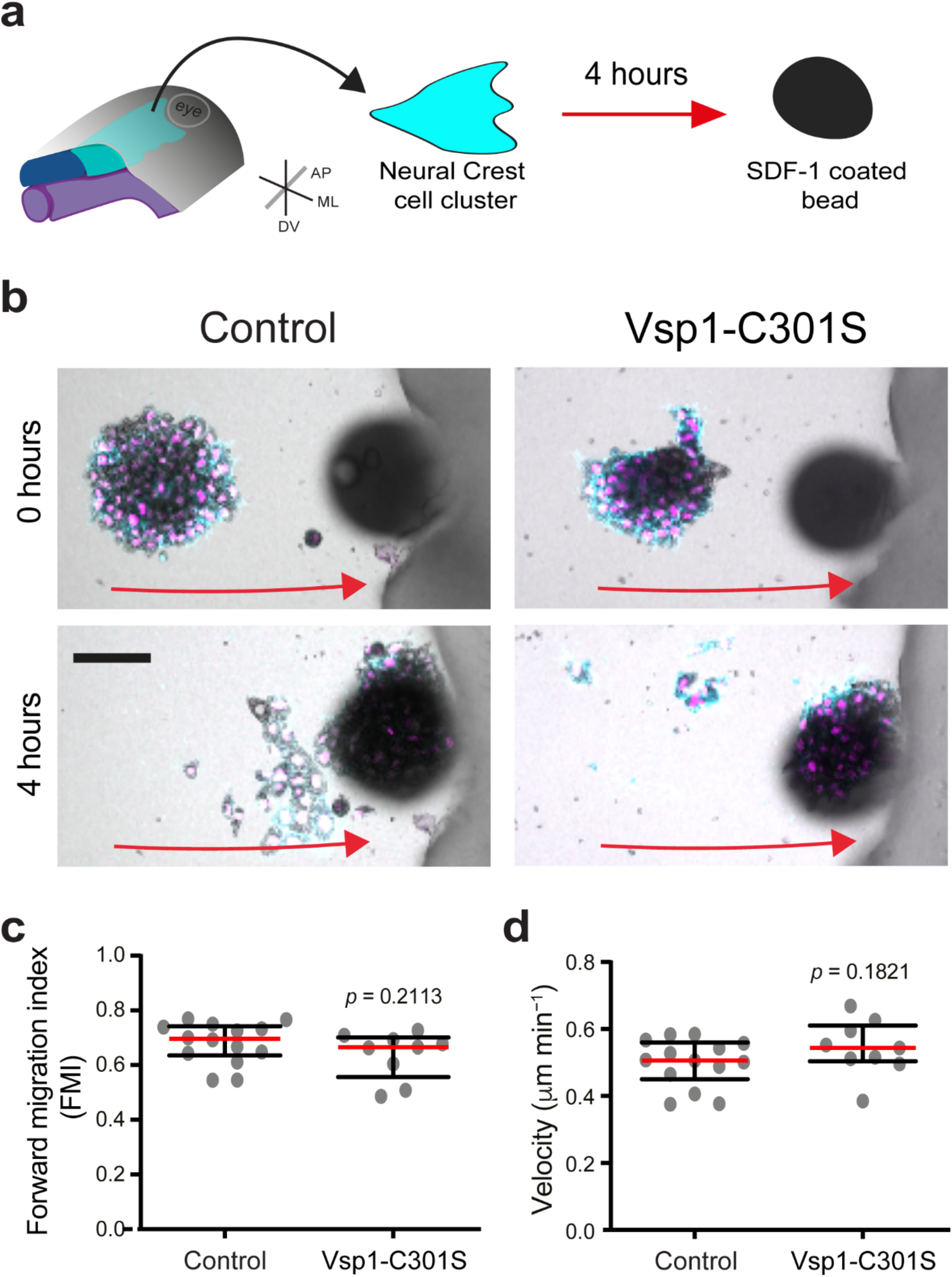
**Vsp1-C301S does not affect chemotaxis of neural crest cells *ex vivo*. a**, Scheme of a typical chemotaxis assay. Neural crest clusters were exposed to a gradient of the chemoattractant SDF-1. **b**, Neural crest clusters migrating towards a SDF-1-coated bead, upper panels show clusters after 5 min of imaging and lower panels show clusters arriving to the SDF-1 bead after 4 hours of time-lapse (red arrows depict the direction of movement). Scale bar, 100 μm. Quantifications of forward migration index (FMI) (**c**) and velocity (**d**) of neural crest clusters, red lines represent median and error bars the interquartile range, Student’s *t*-test, *p*_FMI_ = 0.2113 (two-tailed), *p*_Velocity_ = 0.1821 (two-tailed), *n*_Control_ = 14 clusters, *n*_Vsp1-C301S_ = 9 clusters. **b,** Representative examples from at least three independent experiments; CI = 95%.

**Supplementary Fig. 7.**
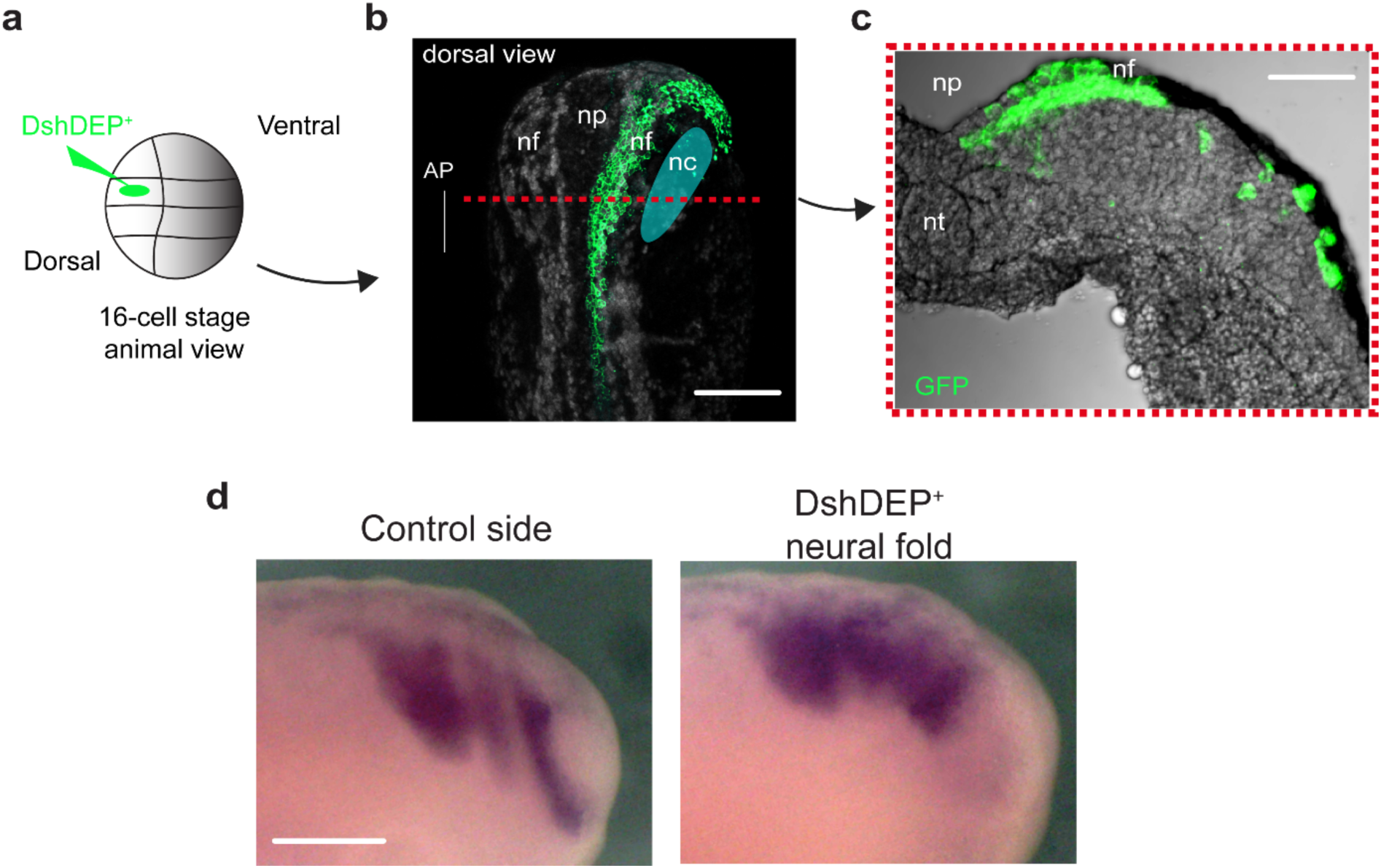
**Targeted injection of DshDEP^+^in the neural fold affects neural crest dCCM in a nonautonomous manner**. **a**, Scheme of neural fold-targeted injection. **b**, Neural fold-tagged embryo at st17 (dorsal view), red dashed line shows the transverse plane of a cryosection presented in **c**. **c**, Cryosection showing the precise distribution of DshDEP^+^and the membrane GFP in the ectoderm at the level of the neural fold. Faded cyan ellipse, anatomical position of the neural crest (nc). nf, neural fold; np, neural plate; nt, neural tube; AP, anteroposterior. **d**, *In situ* hybridisations showing lateral views of embryos hybridised with *sox*8 (a marker of neural crest migration). Scale bars, 200 μm. **b–d**, Representative examples from at least three independent experiments; CI = 95%.

**Supplementary Fig. 8.**
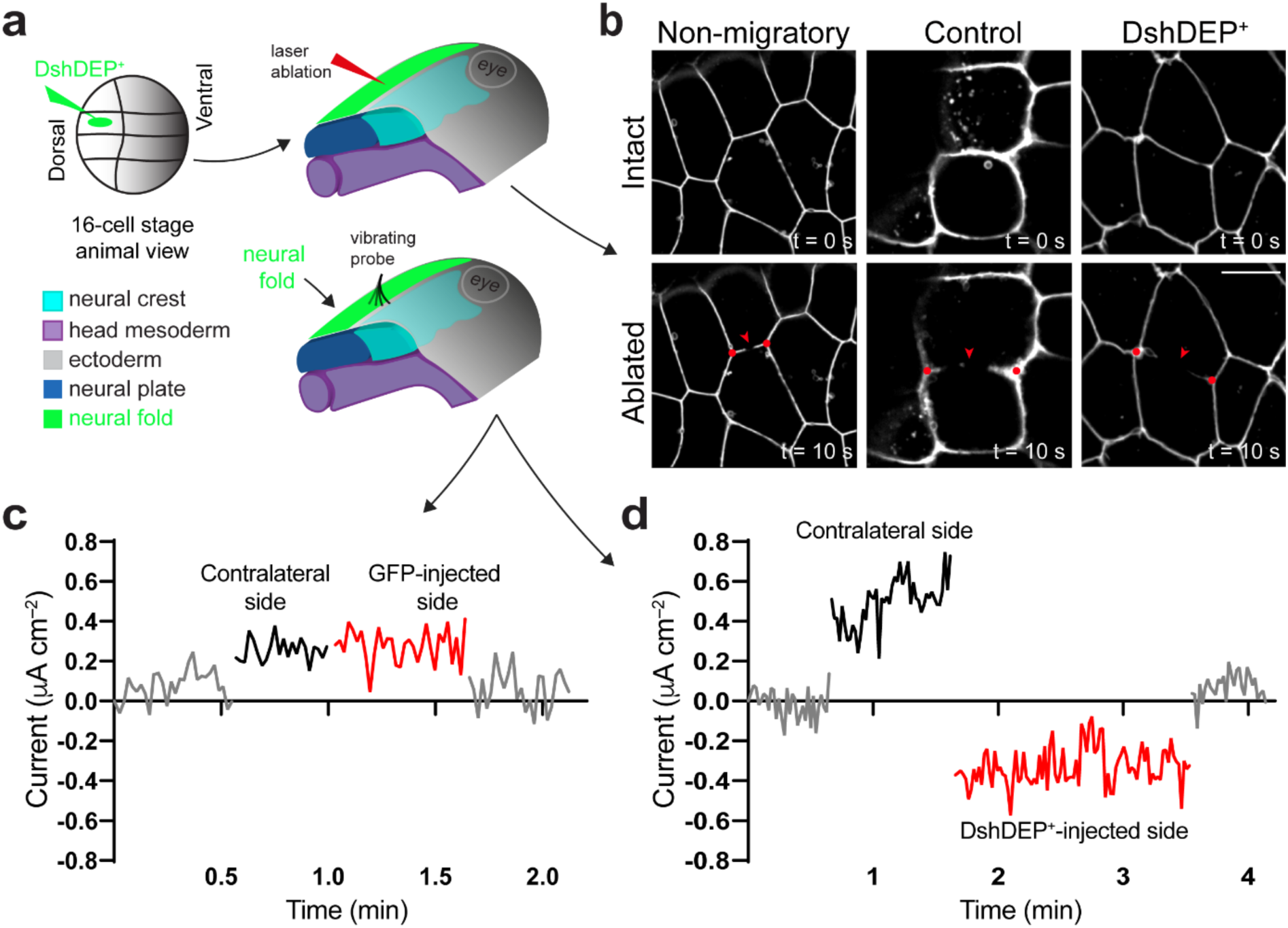
**DshDEP^+^targeted injection in the neural fold reduces membrane tension and electric currents *in vivo*. a**, Scheme of DshDEP^+^targeted injection into the neural fold for laser ablation and current measurements. **b**, Impact of stage and PCP inhibition in neural fold membrane recoil velocity as a function of stored tension (see **Supplementary Video 9**). Injection of GFP does not affect outward currents (**c**) but DshDEP^+^does (**d**). **b**, Representative examples from at least three independent experiments; CI = 95%.

**Supplementary Table 1.**
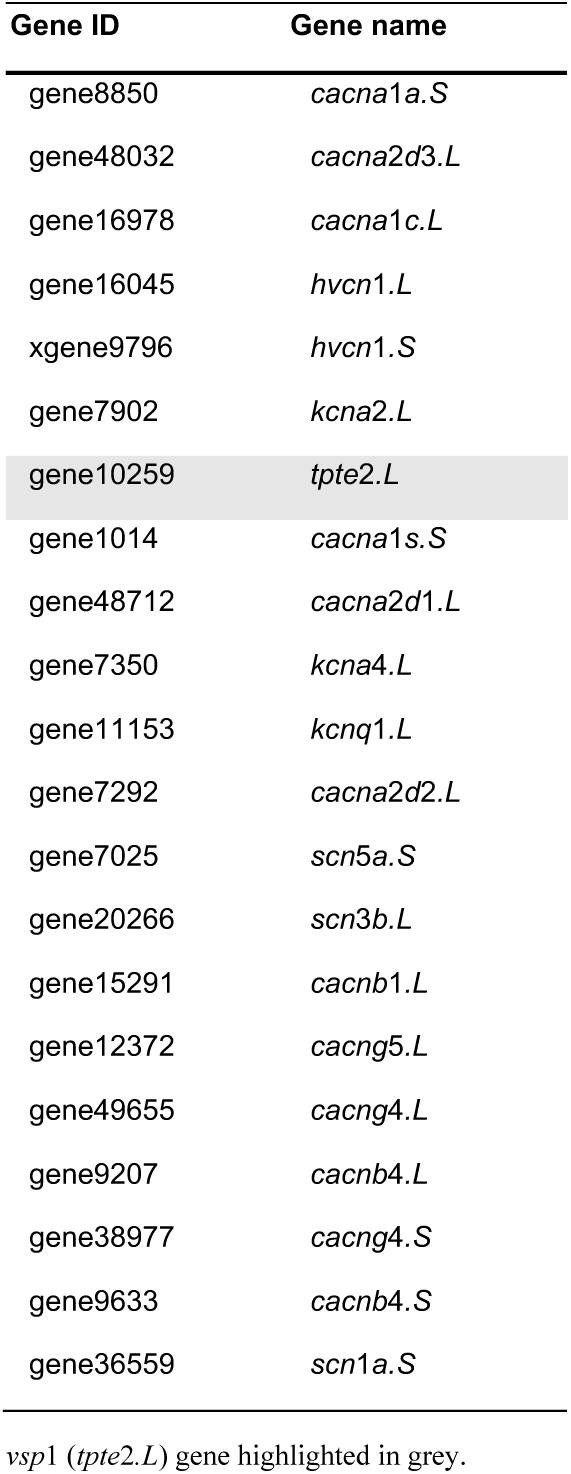
***vsp*1 (*tpte*2*.L*) is present in neural crest cells RNA-seq libraries.** The list provides the ID and names for the most represented ion channels found in our RNA library. These channels were selected as they are members of the voltage-gated ion channel families (https://www.genenames.org). The voltage-sensitive phosphatase *vsp*1 (*tpte*2*.L*) emerged from this list and its paralogous *vsp*2 (*tpte*2*.S*) was not detected, as previously described for these stages^62^. Genes are shown in the same order as in the normalized TMM (Trimmed Mean of M-values) of the RNA-seq data analysis (**Methods**).

**Supplementary Table 2.** ***VSP* (*TPTE*2) expression in cell lines that have been reported to electrotax.** *In silico* analysis reveals expression of orthologous *VSP* in wound healing and cancer cells that are well known to follow the electric fields *in vitro*.

| Cell type | Primary culture or cell line | *VSP (TPTE2)* expression | Electrotactic response |
| --- | --- | --- | --- |
| Mammary epithelium | MCF-10A | Lusche *et al.*, 2018<sup>63</sup> | Lalli and Asthagiri, 2015<sup>64</sup> |
| Fibroblasts | NIH 3T3* | BioGPS (Mouse Cell Type and Tissue Gene Expression Profiles)<sup>65-68</sup> | Huang *et al.*, 2013<sup>69</sup> |
| Human breast cancer | MCF-7* | Expression Atlas (www.ebi.ac.uk/gxa)<sup>70</sup> | Pu *et al.*, 2007<sup>71</sup> |
|  | MDA-MB-231* | Harmonizome (Cell Line Gene Expression Profiles)<sup>67,72</sup> | Wu *et al.*, 2013<sup>73</sup> |
| Macrophages | Primary culture* | BioGPS (Mouse Cell Type and Tissue Gene Expression Profiles)<sup>65-68</sup> | Hoare *et al.*, 2014<sup>74</sup> |

\**VSP (TPTE2)* expression reported in publicly available databases.

## Supplementary Video Legends

**Supplementary Video 1 | Neural crest cells migrate towards the anode of electric fields *ex vivo*.** Representative time-lapses of neural crest clusters in the absence (upper panel) or presence (lower panel) of electric fields (EFs) for 8 h. The lower panel cluster was exposed to EFs of 100 mV mm^−1^ for 4 h, with polarity reversal for another 4 h. Time-lapse setting was 1 picture every 5 min; 96 frames are shown at 7 frames per second. Note the cluster spreading and dispersing in the absence of EFs and the consistent anodal migration in the presence of EFs. Neural crest cells tagged with nuclear RFP (magenta) and membrane GFP (cyan) constructs. Scale bar, 100 μm. Related to Fig. 1.

**Supplementary Video 2 | Neural crest cells electrotaxis *ex vivo* under a range of electric fields recorded *in vivo*.** Representative time-lapses of neural crest clusters exposed to electric fields of 10, 25 and 100 mV mm^−1^. Time-lapse setting was 1 picture every 5 min; 48 frames are shown at 7 frames per second. Note the anodal migration. Neural crest cells tagged with nuclear RFP (magenta) and membrane GFP (cyan) constructs. Scale bar, 100 μm. Related to Supplementary Fig. 2.

**Supplementary Video 3 | Neural crest cells electrotact as a collective but not as single cells *ex vivo*.** Representative time-lapses of a neural crest cluster and an isolated neural crest cell exposed to electric fields of 100 mV mm^−1^. Time-lapse setting was 1 picture every 5 min; 12 frames are shown at 7 frames per second. Note the anodal migration of the cluster and the random migration of the cell. Neural crest cells tagged with nuclear RFP (magenta) and membrane GFP (cyan) constructs. Scale bar, 50 μm. Related to Supplementary Fig. 3.

**Supplementary Video 4 | Vsp1-C301S does not affect neural crest cells motility *ex vivo*.** Representative time-lapses showing the migration of isolated control or Vsp1-C301S-injected cells. Time-lapse setting was 1 picture every 5 min; 12 frames are shown at 7 frames per second. Note the similar motility of both conditions. Neural crest cells tagged with nuclear RFP (magenta) and membrane GFP (cyan) constructs. Scale bar, 30 μm. Related to Supplementary Fig. 5.

**Supplementary Video 5 | Vsp1-C301S impairs neural crest electrotaxis *ex vivo*.** Representative time-lapses of control and Vsp1-C301S-injected neural crest clusters exposed to electric fields of 100 mV mm^−1^. Time-lapse setting was 1 picture every 5 min; 36 frames are shown at 7 frames per second. Note the anodal migration of the control cluster and the random migration of the Vsp1-C301S cluster. Neural crest cells tagged with nuclear RFP (magenta) and membrane GFP (cyan) constructs. Scale bar, 100 μm. Related to Fig. 2.

**Supplementary Video 6 | Vsp1-R152Q allows for neural crest electrotaxis under suboptimal electric fields *ex vivo*.** Representative time-lapses of control and Vsp1-R152Q-GFP-injected neural crest clusters exposed to suboptimal electric fields of 5 mV mm^−1^. Time-lapse setting was 1 picture every 5 min; 72 frames are shown at 7 frames per second. Note the random migration of the control cluster and the anodal migration of the Vsp1-R152Q cluster. Neural crest cells tagged with nuclear RFP construct in cyan; fusion GFP of the Vsp1-R152Q-GFP construct in green. Scale bar, 100 μm. Related to Fig. 2.

**Supplementary Video 7 | Vsp1-C301S affects neural crest directionality *in vivo*.** Representative time-lapses showing lateral views of wild-type embryos in which control or Vsp1-C301S-injected neural crest explants were grafted. Time-lapse setting was 1 picture every 5 min; 72 frames are shown at 7 frames per second. Note the persistent migration of control neural crests in contrast to the random migration of Vsp1-C301S clusters. Neural crest cells tagged with nuclear RFP construct in cyan. Embryo and eye (ellipse) outlined in solid white lines. Scale bar, 200 μm. Related to Fig. 2 and Supplementary Fig. 5.

**Supplementary Video 8 | Vsp1-C301S does not alter neural crest chemotaxis towards SDF-1 *ex vivo*.** Representative time-lapses of control and Vsp1-C301S-injected neural crest clusters exposed to SDF-1-coated beads. Time-lapse setting was 1 picture every 5 min; 48 frames are shown at 7 frames per second. Note the similar directional migration (towards the bead) of both conditions. Neural crest cells tagged with nuclear RFP (magenta) and membrane GFP (cyan) constructs. Scale bar, 100 μm. Related to Supplementary Fig. 6.

**Supplementary Video 9 | Non-migratory and DshDEP^+^-injected neural folds reduce membrane tension *in vivo*.** Representative time-lapses of laser ablations (solid red line) in the membranes of non-migratory (st13), control (pre-migratory st17) and DshDEP^+^ (pre-migratory st17) injected neural folds. Time-lapse setting was 1 picture every 2 seconds; 6 frames are shown at 4 frames per second. Note that the membrane junction recoil in control neural fold is higher than in non-migratory and DshDEP^+^ neural folds. Neural fold cells tagged with membrane GFP construct in grey. Scale bar, 15 μm. Related to Fig. 3 and Supplementary Fig. 8.

**Supplementary Video 10 | PCP in the neural fold controls the directionality of neural crest cells *in vivo*.** Representative time-lapses showing lateral views of embryos in which wild-type neural crest explants (nuclear RFP in cyan) were grafted into control embryos (injected with membrane GFP in the neural fold, shown in grey) or into embryos in which DshDEP^+^ injection was targeted to the neural fold (in grey). Time-lapse setting was 1 picture every 5 min; 60 frames are shown at 7 frames per second. Note that the wild-type neural crest directionally migrates in GFP but not in DshDEP^+^ hosts with targeted neural folds. Scale bar, 200 μm. Related to Fig. 3.

## Notes

### Competing Interest Statement

The authors have declared no competing interest.

